# Porphyrin overdrive rewires pan-cancer cell metabolism

**DOI:** 10.1101/2022.02.18.481061

**Authors:** Swamy R. Adapa, Gregory A. Hunter, Narmin E. Amin, Christopher Marinescu, Andrew Borsky, Elizabeth M. Sagatys, Said M. Sebti, Gary W. Reuther, Gloria C. Ferreira, Rays H.Y. Jiang

## Abstract

**Porphyrin overdrive rewires pan-cancer cell metabolism:** All cancer cells reprogram metabolism to support aberrant growth. Here, we report that cancer cells employ and depend on imbalanced and dynamic heme metabolic pathways for their oncogenic growth. We coined this essential metabolic rewiring ‘porphyrin overdrive’ and determined that it is cancer-universal, cancer-essential, and cancer-specific. While porphyrin overdrive is absent in differentiated cells or somatic stem cells, it is present in patient-derived tumor progenitor cells, demonstrated by single cell RNAseq, and in early embryogenesis. Among the major drivers are proteins involved in biosynthesis of heme intermediates and heme trafficking. CRISPR/Cas9 editing to engineer leukemia cells with impaired heme biosynthetic steps confirmed our whole genomic data analyses that porphyrin overdrive is linked to oncogenic states and cellular differentiation. In conclusion, we identified a dependence of cancer cells on non-homeostatic heme metabolism, and we targeted this cancer metabolic vulnerability with a novel “bait-and-kill” strategy to eradicate malignant cells.

**One-Sentence Summary:** Porphyrin overdrive reprograms cancer cellular metabolism.

## Introduction

Heme biosynthesis is one of the most efficient metabolic pathways in humans. A total of 270 million molecules of heme are produced for every red blood cell (RBC) (D’Alessandro et al., 2017), at a rate of over 2 million RBCs per second (Dean, 2005). Heme, a porphyrin ring encaging most of the iron in humans, functions as an essential prosthetic group of numerous proteins with roles ranging from signal sensing, DNA binding, microRNA splicing and processing to enzymatic catalysis (Ponka et al., 2014). To ensure the enormous output of heme biosynthesis, the supply of substrates, intermediates and end-products in the pathway is tightly regulated and precisely balanced (Hunter and Ferreira, 2011). Heme is a double-edged sword for cell growth; it is essential in the “right amount” (Cao and Dixon, 2016) and if not can be toxic (Malik and Djaldetti, 1980) via unique forms of cell death.

Cancer cells reprogram metabolic pathways to fuel anabolic growth, rapid propagation, and efficient nutrient acquisition(Chaffer and Weinberg, 2011; Martinez-Outschoorn et al., 2017). Although some studies have implicated heme in carcinogenesis through cytotoxic heme-derived compounds (Malik and Djaldetti, 1980), lipid peroxidation (Martin et al., 2018), oxidative damage (Sohoni et al., 2019), intestinal flora toxicity (Ijssennagger et al., 2015) and energy production (Fiorito et al., 2021), little is known about how heme biosynthesis deregulation and heme trafficking alterations contribute to tumor dependence on heme and its biosynthetic pathway for survival. Here we describe ‘porphyrin overdrive’, a novel cancer cell metabolic reprogramming characterized by imbalanced and dynamic heme metabolic pathways essential for cancer cell growth.

“The Warburg effect” (Vander Heiden et al., 2009) and “glutamine addiction” (Shelton et al., 2010) are two altered forms of metabolism found in cancer cells. Therapeutic intervention based on these altered cancer metabolic pathways are challenging because glycolysis and glutamine metabolism are required in every cell and thus such therapeutic approaches lead to non-specific toxicity. By contrast, we propose that porphyrin overdrive is an ideal cancer metabolic pathway for therapeutic targeting as we demonstrate that it is: 1) cancer specific (*i.e.*, it is absent in normal cells), 2) universal (*i.e.*, it is present in all cancers), and 3) cancer cell essential (*i.e.,* cancer cells require it for survival).

## Results and Discussion

### Cancer is characterized by aberrant heme metabolism

Compared to normal cells, cancer cells have enhanced metabolic dependencies (Levine and Puzio-Kuter, 2010; Luengo et al., 2017; Martinez-Outschoorn et al., 2017). Our initial data analyses of genome-scale CRISPR/Cas9-gene loss-of-function from the publicly available project DepMap screens of cell lines derived from metastatic cancers (Meyers et al., 2017) indicated that cancer cells depend on heme synthesis (Fig. S1) (Supplementary Table S1). Significantly, these metastatic cancers developed dependencies on heme metabolism-related proteins, such as uroporphyrinogen III decarboxylase (UROD), the enzyme that catalyzes the fifth step in the heme biosynthetic pathway. These cancers also depend on a set of hemoproteins, such as cytochrome c-1 (CYC1), succinate dehydrogenase complex subunit C (SDHC) for cellular respiration, and DGCR8 for microRNA biogenesis(Nguyen et al., 2015), each of which uses heme as a co-factor. Unexpectedly, the *FECH* gene encoding ferrochelatase that catalyzes the final critical step of heme biosynthesis is dispensable in several cancers, suggesting that cancer cell lines are capable of bypassing endogenous biosynthesis of heme *in vitro*, yet are still dependent on genes encoding enzymes that mediate intermediate steps of heme biosynthesis. This initial analysis suggests that cancer is dependent on imbalanced heme metabolism (Fig. 1A).

**Figure 1.**
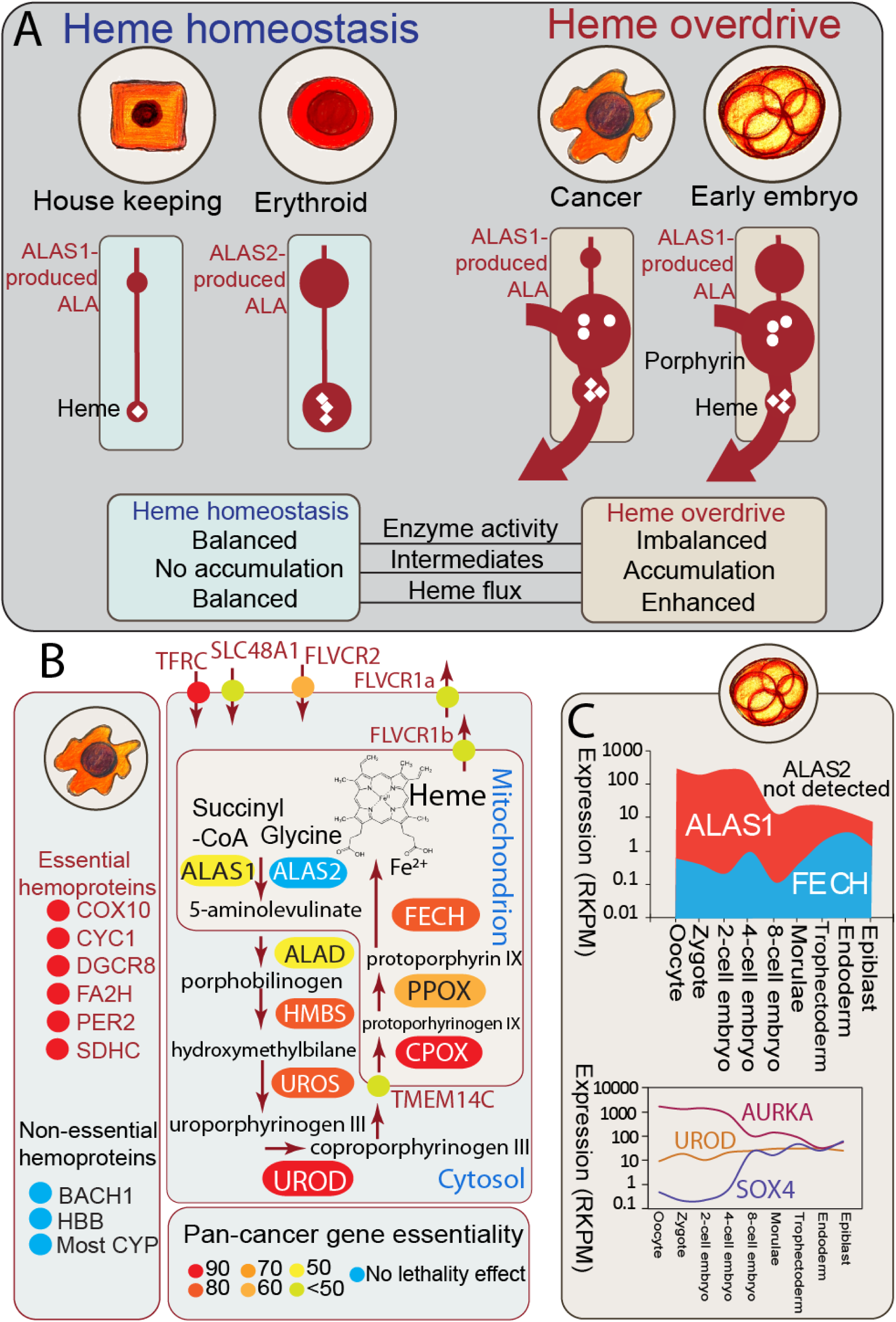
Defining porphyrin overdrive with CRISPR data and early human embryonic stem cell single cell RNAseq. (**A**) **Schematic illustration of heme homeostasis and porphyrin overdrive.** Heme homeostasis refers to the heme biosynthetic pathway in normal cells, with the housekeeping enzyme ALAS1 catalyzing the first and rate-limiting step in all cells, except precursor erythroid cells, where the erythroid-specific ALAS isozyme ALAS2 catalyzes the rate-determining step. Porphyrin overdrive refers to the imbalanced (i.e., with aberrantly increased and suppressed enzyme activities) pathway in cancer cells and early human embryonic stem cells, with the ALAS1 isozyme controlling ALA production regardless of the cell type. White circles and white diamonds represent porphyrin intermediates and heme, respectively; Erythroid refers to erythroid precursor cell. Imbalanced enzyme activities in porphyrin overdrive are indicated by different size of product pools (dark red circle sizes). ALA, 5-aminolevulinate; ALAS, ALA synthase. (**B**) **Defining porphyrin overdrive with CRISPR KO-derived essentiality data in over 300 cancer cell lines.** Analysis of CRISPR essentiality data enables mapping of porphyrin overdrive in 27 major types of cancer. Color mapping indicates average cancer cell growth dependence revealed by CRISPR KO of a given gene. UROD has the highest pan-cancer essentiality. (**C**) Early human embryonic stem cells show features of porphyrin overdrive. The expression levels for *ALAS1* and *FECH* differ over 100-fold during the first rounds of embryonic stem cell divisions, but they normalize with the early embryonic development progression. AURKA and SOX4 show expected patterns of expression dynamics during embryogenesis. [Abbreviations: ALA, 5-aminolevulinate; ALAS1, housekeeping ALA synthase; ALAS2, erythroid-specific ALA synthase; ALAD, ALA dehydratase (aka porphobilinogen synthase); BACH, transcription regulator BACH (BTB and CNC homology); CPOX, coproporphyrinogen oxidase; COX10, cytochrome C oxidase assembly homolog 10; CYC1, cytochrome C1; CYP, cytochrome P450 family; DGCR8, DGCR8 microprocessor complex subunit; FAH2, fatty acid 2-hydroxylase 2; FECH, ferrochelatase; FLVCR, feline leukemia virus subgroup C receptor family; FLVCR1a, FLVCR member 1a; FLVCR1b, FLVCR member 1b; FLVCR2, FLVCR member 2; HBB, hemoglobin β-subunit; HMBS, hydroxymethylbilane synthase; PER2, period circadian regulator 2; PPOX, protoporphyrinogen oxidase; RKPM, reads per kilobase million; SDHC, succinate dehydrogenase complex subunit C; TFRC, transferrin receptor; TMEM14C, transmembrane protein 14C; UROD, uroporphyrinogen decarboxylase; UROS, uroporphyrinogen III synthase].

Next, we investigated the genes that encode proteins acting as gatekeepers for heme biosynthesis. The first committed and key regulatory step in mammalian heme biosynthesis is catalyzed by 5-aminolevulinic acid (ALA) synthase (ALAS; EC 2.3.1.37) (Ponka et al., 2014). Two chromosomally distinct genes, *ALAS1* and *ALAS2*, encode the housekeeping and erythroid-specific ALAS isoforms, respectively (Ponka et al., 2014). While *ALAS1* is expressed in every cell, the expression of *ALAS2* is restricted to developing erythrocytes (Ponka et al., 2014). We examined the data generated from genome-wide CRISPR-based screens of diverse cancer cell lines, which were designed to provide a complete gene essentiality data set for these cell lines (Meyers et al., 2017; Tsherniak et al., 2017). In contrast to *ALAS2*, essentiality of *ALAS1* is a feature of diverse cancer cells independent of cancer type (Fig. S2A). Consistent with these forward genetics results, the gene expression patterns in ∼10,000 patient tumors from the Genotype-Tissue Expression (GTEx) (Ardlie; Lonsdale et al., 2013) and The Cancer Genome Atlas (TCGA) (Chang et al., 2013) datasets show that *ALAS2* expression is absent in most tumors except myeloid leukemias (Fig. S2B). These results show that, despite ALAS2 being responsible for most heme production (∼85%) in humans, many cancer cells, including erythroleukemia of red blood cell lineages (*e.g.*, HEL), require ALAS1.

### Porphyrin overdrive operates in diverse cancers and early embryos

To better understand the molecular and genetic mechanisms underlying the cancer dependency on heme metabolism, we determined gene essentiality from genome-scale CRISPR/Cas9 loss-of-function screens in over 300 human cancer cell lines covering different cell lineages and estimated gene dependency (Meyers et al., 2017; Tsherniak et al., 2017). Then, we focused on the gene dependencies associated with the eight enzymatic steps of the heme biosynthetic pathway by computing the respective pan-cancer essentiality scores, similar to the lethality scores used in the development of a cancer dependency map (DepMap) (Dempster et al., 2019; Pacini et al., 2021). Gene essentiality score values lower than zero indicate decreased cell growth/viability with loss of gene X, and hence the lower the X gene essentiality score, the more dependent are cells on the X gene. Consistent with recent cross-platform studies (Dempster et al., 2019; Pacini et al., 2021), over 2000 genes were defined as pan-cancer essential, based on the distribution of their essentiality scores in over 300 cancer lines derived from 27 of the most prevalent cancer types. The gene essentiality scores were confirmed by using the same method on the total DepMap collection (DepMap release 21Q3) of over 1000 cell lines (Pearson’s R > 0.9, p < 0.001). Strikingly, instead of finding the expected complete and balanced heme biosynthetic pathway characteristic of normal cells, we found that the survival of cancer cells depends on the various steps of the heme biosynthetic pathway in a “non-balanced” manner. In hundreds of different types of cancers, the tumor cells particularly rely on the intermediate enzymatic steps of heme biosynthesis, as assessed by loss of cellular viability upon CRISPR/Cas9-mediated gene ablation in whole genome loss-of-function screens (Fig. S3) (Supplementary Tables S2 and S3). Specifically, as shown in Fig. 1B and Fig. S3, the pan-cancer gene essentiality is the highest for the genes encoding the enzymes responsible for the fifth and sixth steps of the pathway, UROD and coproporphyrinogen III oxidase (CPOX), respectively. This result was notably surprising, as the genes for the ALAS isoforms, which catalyze the initial and rate-limiting step in the pathway, and the gene for FECH, the final enzyme in the pathway that catalyzes heme production, had lower essentiality values and were dispensable in several cancers (Fig. 1B and S3). Though the heme biosynthesis genetic dependences of 27 major forms of human cancer vary, the estimated patterns of essentiality across the cancer lines clearly revealed a range of heme and heme precursor requirements. These results uncovered the “unbalanced” nature of heme biosynthesis in cancer, with many cancers surviving without functional first and terminal enzymatic steps but depending on the intermediate steps (Fig. S3 and Fig. S4). The gene essentiality analysis also highlighted the importance of heme trafficking (e.g., heme importer FLVCR2) and key hemoproteins (*e.g*., CYC1) in cancer cells survival (Fig. 1B and Fig. S3).

To examine cancer heme metabolism *in vivo*, we analyzed the *in vivo* mouse CRISPR/CAS9 loss-of-function and metabolic essentiality data from pancreatic and lung cancer models (Zhu et al., 2021). The genetic screen (Zhu et al., 2021) indicates that most metabolic gene essentialities were similar between the *in vitro* and *in vivo* settings, except for the heightened requirement of the intermediate steps of heme biosynthesis *in vivo*. Strikingly, out of the ∼2900 examined metabolic genes(Zhu et al., 2021), the genetic dependencies of both murine pancreatic and lung cancers were only increased for the enzymes responsible for the intermediate steps of the heme biosynthetic pathway (Fig. S5) (Supplementary Table S4). The essentiality extent of either *ALAS* or *FECH*, which code for the first and terminal enzymes of the heme biosynthesis pathway, respectively, did not differ significantly between the *in vivo* murine cancer models. This analysis of the CRISPR/Cas 9 loss-of-function screens performed in mouse model systems in pancreatic and lung models revealed that aberrant heme metabolism has increased importance in both of these cancer types that were assessed (Fig. S5).

To assess heme biosynthesis in patient tumor samples, we analyzed data available from the GTEx project (Ardlie; Lonsdale et al., 2013) and TCGA program (Chang et al., 2013). We used methods designed for pairwise comparative gene expression of GTEx and TCGA datasets (Tang et al., 2017). In over 80% of all tumors, the genes for hydroxymethylbilane synthase (HMBS), the third enzyme of the heme biosynthetic pathway, and the heme exporter FLVCR1 were among the most upregulated (Fig. S6A-B) (Supplementary Table S5). By the same criteria, the onco-signaling genes *AURKA*, *KRAS* and *MYC* were upregulated in over 90%, 60% and 50% of all tumor types, respectively. In contrast, expression of the genes encoding the second and terminal enzymes of the heme biosynthetic pathway, aminolevulinate dehydratase (ALAD, a.k.a. porphobilinogen synthase) and ferrochelatase (FECH), respectively, are downregulated suggesting the presence of a buildup of intermediate heme precursors in tumors. *HPX*, the gene for the heme-binding and scavenger hemopexin (Tolosano et al., 2010), was also downregulated in most of the tumors (Fig. S6C). This finding, which corroborates the previously reported role of hemopexin as a key player for the checkpoint in cancer growth and metastases (Canesin et al., 2020), leads compared the gene expression patterns in ∼10,000 tumors *vs*. those in normal tissues us to suggest that heme content is high in the tumor microenvironment. *ALAS2* belongs to the 10% of genes that were not expressed in most of the large tumor collections, except myeloid leukemias (<2% of total tumors). Overall, our comparative analysis of the differential gene expression in tumor tissues, spanning 31 cancer types, indicated that only a few of the nine genes encoding the enzymes of the heme biosynthetic pathway were upregulated in cancer cells.

Based on the gene essentiality results from both *in vitro* (Fig. 1B, Fig. S3 and Fig. S4) and *in vivo* (Fig. S5) settings, the pan-cancer gene expression patterns (Fig. S6), the detected porphyrin accumulation in a wide range of tumors (Fiorito et al., 2020), and the reported elevated heme flux in diverse cancer cells (Hanna et al., 2016), we defined the salient porphyrin overdrive-associated features. They are: 1) heme biosynthesis enzymes with aberrantly increased (HMBS, UROS and UROD) and suppressed (ALAD and FECH) activities, 2) accumulation of heme precursors (porphyrins), and 3) enhanced heme flux (Fig. 1, A and B). This cellular status contrasts with the tightly regulated steady-state or homeostasis, defined by a flawless enzyme-catalyzed channeling of substrates to products along the heme biosynthetic pathway and circumventing the toxicity and instability of the pathway intermediates, while ensuring that the cell heme requirements are met (Ferreira, 2013). With porphyrin overdrive, the production of heme precursor molecules exceeds that of heme, porphyrin intermediates accumulate, and “imbalanced” heme biosynthesis of increased porphyrin intermediates and decreased end-product arises. Likely, to compensate for the reduced amount of heme synthesized, heme flux is increased(Fiorito et al., 2020). Cancer cells appear to have adopted a dependence on this seemingly “inefficient” metabolic pathway designed to produce aberrant levels of heme intermediates with unknown biological functions to date.

Early embryonic stem cells represent a special metabolic state that bears similarities with that of cancer cells (Smith and Sturmey, 2013). Therefore, we also studied the expression of the heme biosynthetic enzyme-encoding genes in human preimplantation embryos and embryonic stem cells. Specifically, we explored a high-resolution data set generated upon single-cell RNA-sequencing in human early embryonic cells and embryonic stem cells (Li et al., 2017; Yan et al., 2013). In the human zygote and early preimplantation embryos, the level of *ALAS1* expression is 100-fold higher than that of *FECH* (Fig. 1C). This large difference in gene expression for two key enzymes in heme biosynthesis was observed in all 17 early human embryos examined (Supplementary Table S6). The large difference in expression levels is restricted to very early embryos because it rapidly narrows, via an increase in *FECH* expression and a decrease in *ALAS1* expression after initial rounds of embryonic cell division. *ALAS2* expression is not detected in any of the early human embryos, which is congruent with the non-essentiality of *ALAS2* in all examined cancers. As a control, the genes *AURKA* and *SOX4* show expected expression patterns during early embryogenesis (Penzo-Méndez et al., 2007; Sasai et al., 2008). The similarity of these gene expression data with those observed in cancer cells supports the possibility that a form of porphyrin overdrive operates in preimplantation human embryos.

## Porphyrin overdrive is absent in normal cells

To evaluate if porphyrin overdrive is specific to cancer cells (and possibly preimplantation embryos), we investigated and compared heme biosynthesis in diverse types of non-cancerous cells, *i.e.,* normal differentiated cells, replicating fibroblasts, human primary hepatocytes, and human primary hematopoietic stem cells. We looked for evidence of porphyrin overdrive, defined by a non-homeostatic, imbalanced heme biosynthetic pathways and heightened accumulation of heme pathway intermediates. Using published datasets, we found that porphyrin overdrive, is absent in normal differentiated cells and somatic stem cells (Fig. 2A) First and consistent with this finding, normal human peripheral blood mononuclear cells (PBMCs) do not accumulate heme intermediates even upon ALA-induction (Oka et al., 2018). Second, colon cancer cells incubated with ALA to induce protoporphyrin IX (PPIX) production do accumulate PPIX, while no significant PPIX accumulation is observed in replicating human colon-derived fibroblasts (stromal cells) previously incubated with ALA (Krieg et al., 2002). Further, the increased HMBS activity in colon cancer cells relative to stromal cells (Krieg et al., 2002) is consistent with a buildup of heme intermediates including PPIX. Third, using our previously described primary human hepatocyte system (Maher et al., 2020), we demonstrate that normal, metabolically active, primary hepatocytes do not accumulate porphyrins as indicated by the absence of the PPIX fluorescence (Fig. 2B). By contrast, >99% of liver cancer cells produced and accrued PPIX upon induction with ALA, demonstrating a canonical features of porphyrin overdrive.

**Figure 2.**
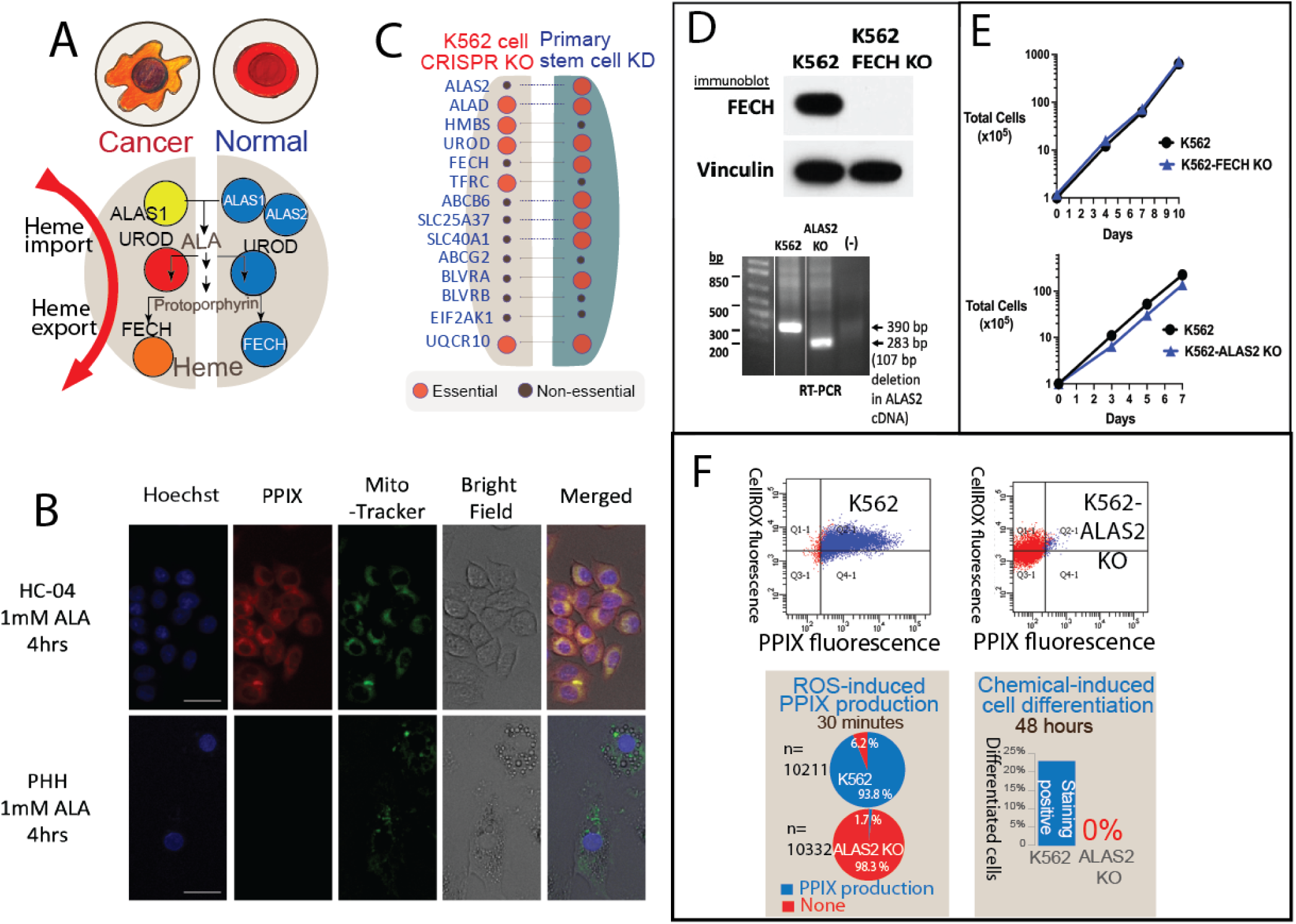
Porphyrin overdrive is unique to cancer cells and absent in normal cells. (**A**) Schematic illustrations of the key differences of heme metabolism in cancer *vs.* normal cells. Red, orange, and yellow shadings represent cancer gene essentiality as in Figure 1B. Blue colors refer to the normal heme biosynthesis process (**B**) PPIX autofluorescence (red) is detected in HC-04 liver cancer cells but not in primary human hepatocytes (PHH) following ALA addition, indicating PPIX accumulation in cancer cells but not normal cells. Mitotracker green staining indicates viable cells. Over 1000 cells were assessed for each sample (representative cells shown). PPIX red was pan-cancer essentiality analysis (Depmap) (**D**) Validation of K562-FECH KO cell line generated by CRISPR/Cas9 gene editing. Immunoblots of wild-type K562 and K562-FECH KO whole cell lysates show a complete loss of FECH protein in K562-FECH KO cells, with vinculin used as a loading control (top). A 107 bp out of frame CRISPR-induced deletion was identified in the *ALAS2* gene (not shown) and due to in inability to identify a specific ALAS2 antibody for immunoblotting, presence of this deletion was confirmed in RT-PCR analysis of K562-ALAS2 KO cells (bottom). (**E**) Total viable K562 and K562-FECH KO cells (top) and K562 and K562-ALAS2 KO cells (bottom) were determined over time using trypan blue exclusion, and no growth differences were detected. (**F**) ALAS2 KO K562 cells are arrested in an undifferentiable state. K562 cells readily differentiated upon ROS induction (tert-butyl-hydroperoxide) and are committed into erythroid lineage within 48 hrs, as measured by activation of heme production and benzidine stain as a marker of differentiation. In contrast, ALAS2 KO cells do not respond to the ROS inducer (the first step of differentiation) nor commit to erythroid differentiation.

To uncover the role of heme metabolism in normal human stem cells, we analyzed the erythropoiesis data generated by RNAi-based knockdown of gene expression in human hematopoietic progenitor cells (Egan et al., 2015). By mapping normal stem cell gene essentiality following the methods described in Egan *et al*. (Egan et al., 2015), we show that erythropoiesis is heme-dependent in both early progenitor (undifferentiated) cells and differentiated cells from normal human bone marrow-derived hematopoietic stem cells. In contrast to cancer cells, where *ALAS2* is dispensable, normal human hematopoietic stem cells depend on *ALAS2* for survival (Fig. 2C). The differential gene essentiality profiles between cancer and stem cells are consistent with the molecular basis for specific ex-vivo purging of cancer hematopoietic progenitor cells, via ALA-induction/photodynamic therapy as a way to eradicate malignant cells while sparing hematopoietic stem cells (de Lima and Shpall, 2004). Taken together, the distinct genetic essentialities and the markedly different porphyrin accumulation modes between cancer and non-cancerous cells indicate that porphyrin overdrive is absent in somatic stem cells.

### CRISPR/Cas9-mediated knockout of *ALAS2* and *FECH* validate abnormal heme metabolism underlying cell proliferation and oncogenesis

We modified chronic myeloid leukemia K562 cells with CRISPR/Cas9-mediated knockout (KO) of the genes encoding FECH and ALAS2 (Fig. 2D). Under a heme homeostasis model, a cell line without *FECH* should not survive, because *FECH* is a single copy gene in humans and with no functional substitute(Ponka et al., 2014). Indeed, knockdown (KD) of *FECH* in human stem cells led to cell death (Egan et al., 2015). However, under porphyrin overdrive conditions, we predicted that cancer cells rely on heme trafficking and are addicted to the porphyrin intermediates, and thus do not depend on the final step of heme biosynthesis for survival. As hypothesized, K562 cells with the *FECH* edited out (K562-FECH KO) (Fig. 2D) had normal growth (Fig. 2E) under standard culture conditions. The unimpeded growth of the K562-FECH KO cells presumably indicates that at least some cancer cell lines can metabolically function as heme auxotrophs.

As predicted, K562 cells with the *ALAS2* deleted (K562-ALAS2 KO) (Fig. 2D) lost their differentiation capacity although their growth was not hindered (Fig. 2E, F). Given the proposal that oxidation stress, with the involvement of reactive oxygen species (ROS), is the first step in erythroid differentiation of K562 cells (Chenais et al., 2000), we used the organic peroxide tert-butyl hydroperoxide (t-BHP) to trigger ROS-mediated erythroid differentiation. As early as 30 minutes following t-BHP-mediated ROS induction, the parental K562 cells started synthesizing heme, an early step in cellular differentiation (Chenais et al., 2000), while the K562-ALAS2 KO cells failed to initiate this metabolic step, indicated by the lack of heme production (Fig. 2F) (Chenais et al., 2000). Further, at 48 hours, the parental K562 cells readily differentiated into erythroid cells, but the K562-ALAS2 KO cells continued to proliferate and failed to differentiate, as expected because ALAS2 is required for erythroid differentiation. Conceivably, in the absence of endogenous production of heme destined to hemoglobin, the erythroid lineage commitment is blocked, forcing the K562-ALAS2 KO cells to an obligatory porphyrin overdrive path and into a proliferative/non-differentiable, or cancerous, state.

### Single-cell transcriptomics reveals porphyrin overdrive in AML patient biopsies

To study porphyrin overdrive in primary cancer cells, we performed single-cell RNAseq on biopsies from acute myeloid leukemia (AML) patients (Supplementary Tables S7 and S8). For each patient, we obtained around 2000 single-cell transcriptomes, and we analyzed cells of various differentiation stages and cell types. Interestingly, in 2 patients, only a minority of cells (5-10% of total cells) were identified as cancer progenitor cells with hyper proliferation features (Fig. 3A), and they exhibited hybrid features of mature erythroid cells (*e.g*., expressing the genes *HBB* and *GYPA* for hemoglobin β and glycophorin A, respectively) and stem cells (*e.g*., expressing *SOX4*). The cancer progenitor cells (Fig. 3B) had both mature cell features, *e.g.*, *HBB* expression, and early progenitor properties, *e.g.*, proliferative phenotype. Specifically, they were characterized by elevated iron and heme metabolic gene expression, cell proliferation markers, *KRAS* and anabolism process genes, yet they had reduced normal stem cell progenitor marker gene expression *e.g.*, *SOX4* and *JUN* (Fig. 3C) (Merryweather-Clarke et al., 2016). Additionally, cancer/AML progenitor cells exhibited imbalanced heme biosynthesis, inferred, and exemplified by elevated gene expression for enzymes that catalyze the intermediate steps of heme biosynthesis, *e.g.*, HMBS and UROD (Fig. 3D and E). The cancer progenitor cells also had a higher expression level of genes related to heme trafficking such as *FLVCR1* and *DNM2* (Fig. 3C). Our results show that these cancer progenitor cells exhibit features of porphyrin overdrive, consistent with the abnormal heme anabolism and endogenous PPIX accumulation upon ALA administration in leukemia cells compared to normal PBMCs (Oka et al., 2018). Notably, in two solid tumors (Fig. 3F) with independently generated single cell transcriptomes (Tirosh et al., 2016; Wu et al., 2020), the genes encoding the enzymes for the intermediate steps of heme biosynthesis were similarly overexpressed in the cancer progenitor cells (Fig. 3F).

**Figure. 3.**
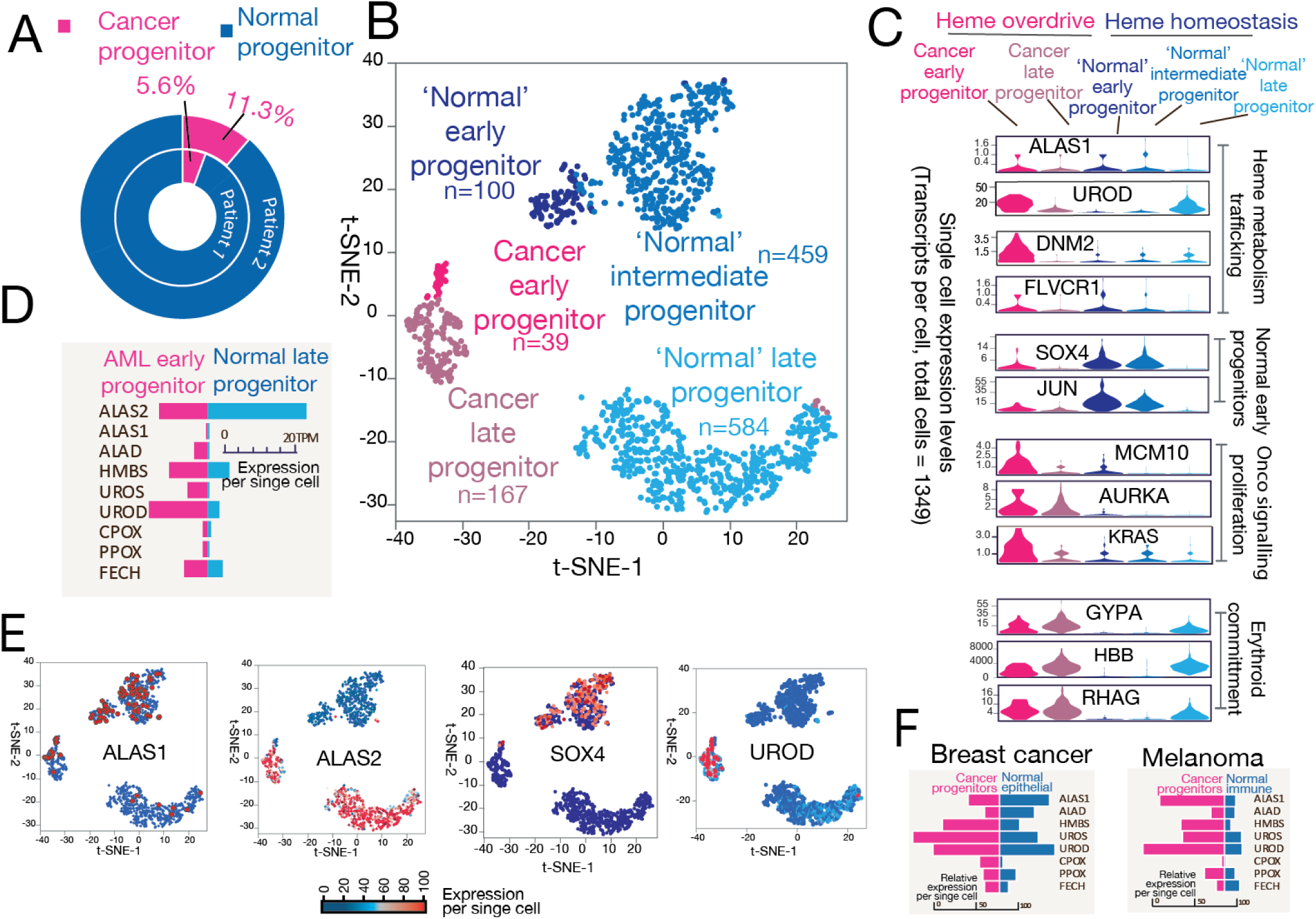
Single-cell RNA sequencing of human AML bone marrow samples supports porphyrin overdrive as a hallmark of cancer cells. Single-cell transcriptomes were obtained from bone marrow biopsies of AML donors with over 60% blasts. (**A**) Single-cell population composition shows only a minor fraction of the cells are cancer progenitors, which are defined by transcriptome embedding and clustering analysis. (**B**) Identification of distinct single cell populations in the AML patient bone marrow samples by t-SNE (t-distributed **S**tochastic **N**eighbor **E**mbedding) analysis. A total of 1349 high-quality transcriptomes from patient biopsies were used. (**C**) Representative genes for proteins associated with heme metabolism, normal early erythroid development, cell proliferation and erythroid commitment process are plotted. (**D**) *UROD*, the gene encoding the fifth enzyme of the heme biosynthetic pathway, is highly expressed in cancer early progenitors. *HMS* and *UROS*, which encode other intermediate step-catalyzing enzymes, are also over expressed in cancer progenitors. (**E**) Marker genes delineate cell populations from the patient samples. Note the complementary expression patterns of *ALAS1 vs*. *ALAS2*. (**F**) Solid tumor single cell RNAseq shows overexpression of the genes for the enzymes responsible for the intermediate heme biosynthetic pathway steps in breast cancer and melanoma (Tirosh et al., 2016; Wu et al., 2020).

From our single cell transcriptome results, we searched for clues of specialized oncogenic microenvironments computationally that could support enhanced substrate trafficking and molecular interactions. Interestingly, only the cancer progenitor cells highly express cell-to-cell interaction-related genes, *e.g.*, *ICAM4*, *MAEA*, *ITGA4* (Fig. S7A), which are critical in establishing the stem cell niche for erythropoiesis (Manwani and Bieker, 2008). We infer that the specific expression of these niche interaction genes suggests that the cancer progenitor cells exist in a specialized niche with intimate contact with neighboring cells and extracellular matrix, and establish evidence for future characterization of this unknown microenvironment.

Expression was remarkably enhanced for genes encoding proteins involved in metabolic substrate trafficking processes in the cancer progenitor cells, including genes for key protein players in lipid import purine and pyrimidine salvage pathways (Fig. S7B). These specific metabolic flux related genes, such as *DNM2* (endocytosis), *APOC1* (lipid transport), *MRI1* (L-methionine salvage), *MTAP* (NAD salvage), and *SLC29A1* (nucleoside import) indicate that the cancer progenitor cells represent hubs of metabolic trafficking activities. In particular, the lipid import gene *APOC1* is expressed in cancer progenitor cells, but not in the other cell populations, including normal progenitor cells, from the same patient samples (Fig. S8, A and B). The metastatin S100 family members linked to tumor niche construction (Lukanidin and Sleeman, 2012) are also enriched in the progenitor cells. These potential dynamic metabolic trafficking patterns inferred from the abundant expression of metabolite import genes in cancer early progenitors (Fig. S7C) are reminiscent of ‘metabolic parasitism’ (Icard and Lincet, 2013), a bioenergetically favorable process that utilizes energy-rich and pre-existing macromolecules to fuel robust cell proliferation.

### Targeting cancer porphyrin overdrive with a bait-and-kill strategy

Two aspects of cancer porphyrin overdrive offer opportunities for therapeutic intervention. First, an imbalanced heme biosynthetic pathway (*i.e.*, with aberrantly increased and suppressed enzyme activities) is absent in normal cells. Second, porphyrin overdrive is inducible hundreds to thousands-fold in cancer cells, but not in non-cancerous cells. In fact, photodynamic therapy (PDT), which uses ALA to induce biosynthesis and accumulation of photosensitizing porphyrins, and exemplifies the inducible principle of porphyrin overdrive, has been used for cancer imaging and treatment for decades (Krammer and Plaetzer, 2008). Thus, porphyrin overdrive may be a unique metabolic reprogramming feature of cancer cells that can be therapeutically exploited. Since an imbalanced heme flux is fundamental to porphyrin overdrive, cancer cells may be made particularly vulnerable to PPIX-induced cytotoxicity. Selective cancer cell death could possibly be achieved by activating PPIX-induced cytotoxicity with a two-step “bait-and-kill” strategy.

Knowing that cancer cells readily take up the heme precursor ALA and bypass the first and rate-limiting step of porphyrin biosynthesis catalyzed by ALAS, we set to assess our engineered two-step process for cancer metabolic death. This two-step process involves first, to ‘bait’ cancer cells with ALA to induce PPIX accumulation, and second, to ‘kill’ the cancer cells with a compound that exploits the metabolic stress associated with porphyrin accumulation. As expected, ALA addition to the culture medium did enhanced PPIX buildup in K562 (Fig. 4A) and HEL (Fig. 4B) cells, as demonstrated by the few hundred-to-thousand-fold PPIX concentration increase in relation to respective cells cultured in the absence of ALA (Fig. 4A and B). Further, HEL cell viability was not affected by supplementing the culture medium with ALA (1 mM final concentration) or ALA and DMSO (1 mM and 0.1% final concentrations, respectively) (Fig. 4C). These results validated that the experimental criteria had been met to proceed with the second step of the “bait-and-kill” strategy.

**Figure 4:**
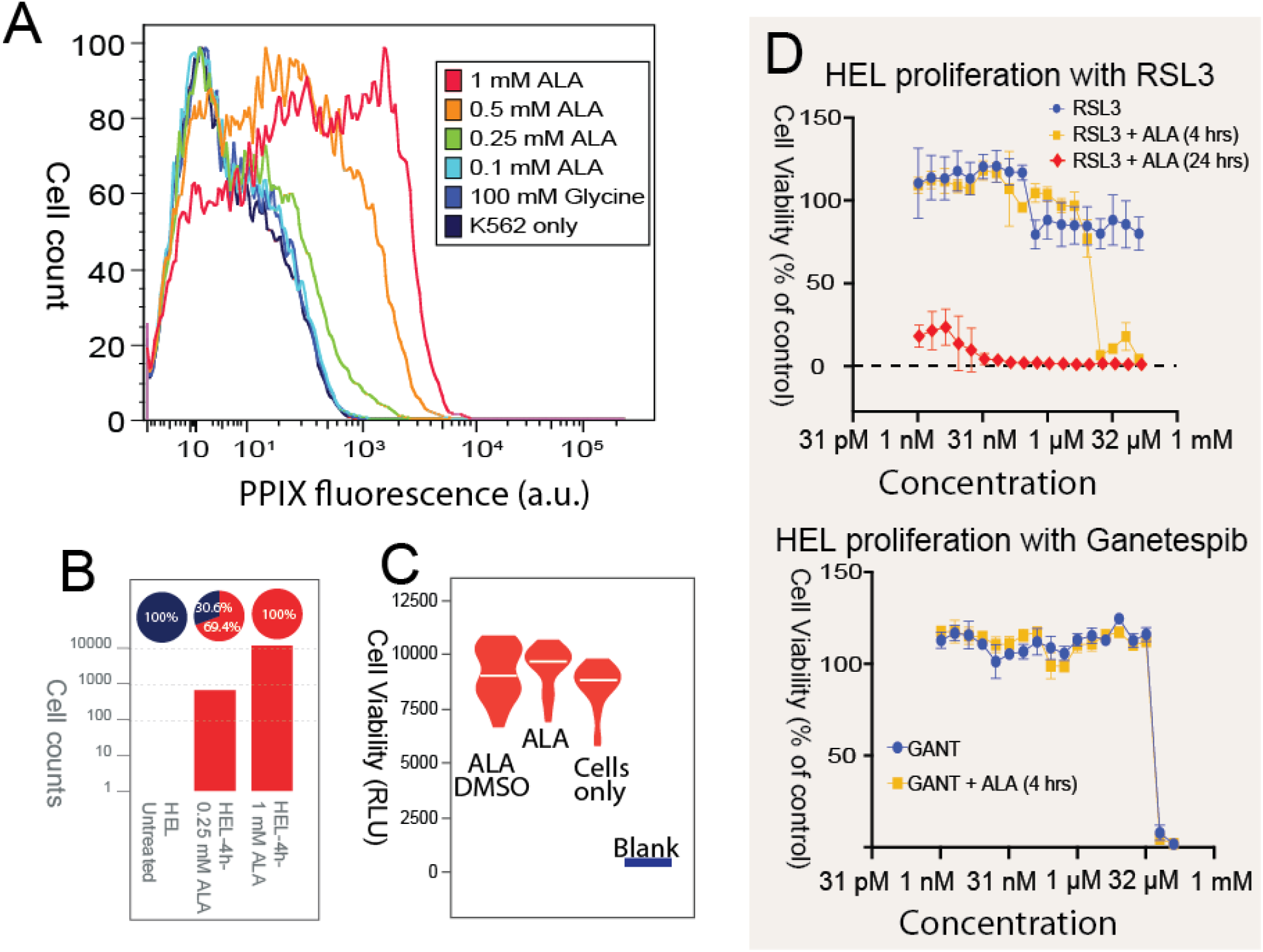
Targeting porphyrin overdrive in cancer cells with a bait-and-kill strategy. (**A**) The concentration of PPIX accumulated in K562 cells depends on the ALA concentration supplemented to the culture medium. K562 cells were cultured in the absence or presence of 100 mM glycine or ALA (0.1 mM, 0.25 mM, 0.5 mM, and 1.0 mM) at 37 °C for 24 h, and PPIX fluorescence was measured by flow cytometry. [a.u., arbitrary units] (**B**) ALA-induced PPIX accumulation in HEL cells. PPIX accumulated in 100% of the HEL cells (n >10,000 cells) as detected by PPIX fluorescence 4 hours after treatment with 1.0 mM ALA. The red sectors of the pie chart above the graph bars indicate the percentage of PPIX-accumulating HEL cells, while those in blue indicate the percentage of cells with no accumulated PPIX. The normalized PPIX values (%) indicated on the pie charts were obtained by dividing the PPIX fluorescence by the PPIX fluorescence intensity value for HEL cells incubated in 1.0 mM ALA-containing medium for 4 h, which was arbitrarily assigned 100%. (**C**) Cell viability is unaffected using ALA (“bait”) as inducer of PPIX accumulation. HEL cells were cultured in the absence or presence of ALA (1 mM) or ALA (1 mM) and DMSO (0.1%) at 37 °C for 24 h, and their viability was calculated from the generated luminescence upon reaction with CellTiter-Glo. The assays were conducted in triplicate. [RLU, relative luminescence units]. (**D**) Dose-dependent response of HEL cell viability for cells treated with increasing concentrations of RSL3 (Top plot) or ganetespib (Bottom plot). Cell viability was determined following incubation with a wide concentration range of RSL3 (top plot) or ganetespib (GANT) (bottom plot) either in the absence or presence of ALA (1.0 mM) for 4 or 24 h. [Note that the concentration of either RSL3 or ganetespib spans a range from pM to 150 mM.]

To exploit the metabolic stress associated with porphyrin accumulation (*e.g.*, oxidative stress, lipid peroxidation), we treated the human erythroleukemia HEL cell line with a range of concentrations of RSL3, an inhibitor of the antioxidant enzyme glutathione peroxidase 4 (GPX4), a phospholipid hydroperoxidase that protects cells against membrane lipid peroxidation. As such, inhibition of GPX4 with RSL3 induces ferroptosis, an iron-dependent form of programmed cell death highly relevant to iron and heme metabolism. Pretreatment of HEL cells with ALA increased their sensitivity towards RSL3 at least 1000-fold (Fig. 4D). Even at a RSL3 concentration of 100 μM, the viability of HEL cells remained similar to that of cells cultured in the absence of the RSL3 inhibitor. However, the viability of HEL cells grown in the presence of ALA for 4 h diminished with RSL3, reaching non-viability at a concentration of 17 μM. The decline was yet more acute when the incubation time of the HEL cells with ALA was extended to 24 h, and the cells became non-viable with 16 nM RSL3. In contrast, ALA pretreatment did not sensitize cells to ganetespib, a heat shock protein inhibitor with antineoplastic activity (Alexandrova et al., 2015), suggesting that ALA is not a general inducer or sensitizer of cell death. Specifically, at ganetespib concentrations greater than 33 μM, HEL cells persist viably regardless of being grown in the presence or absence of ALA (Fig. 4D).

In conclusion, we report that the essentiality pattern of the genes for the enzymes of the heme biosynthetic pathway varies significantly between human healthy and cancer cells. With the genes for the third, fourth and fifth enzymes being particularly critical for cancer cell proliferation, accumulation of porphyrin intermediates exceeds heme production. We propose that cancer cells, in an attempt to counteract unbalanced heme biosynthesis, scavenge heme from their environment and deploy vigorous heme trafficking strategies. We designate the combined non-homeostatic heme synthesis and enhanced heme trafficking as porphyrin overdrive. We suggest that porphyrin accumulation due to porphyrin overdrive and the resulting oxidative stress presents a vulnerability of cancer cells that may be amenable to novel therapeutic strategies to specifically destroy cancer cells.

## Methods

### Cell culture, growth and quantification

K562 (ATCC, Cat No. CCL-243) and HEL (ATCC, Cat No. TIB-180) cells were grown in RPMI 1640 medium supplemented with 10% fetal bovine serum, gentamicin PennStrepNeo and 2 mM L-glutamine at 37 °C in a humidified 5% CO_2_ atmosphere. The cells were plated at 1-2×10^5^/mL and counted by trypan blue (Corning, Cat No. 25-900-CI) exclusion over time.

### CRISPR and erythropoiesis assays

K562 cells containing deletions in *ALAS2* or *FECH* were created using a multi-guide strategy via nucleofection (Lonza) of Cas9/RNP complexes (Gene Knockout Kit v2, Synthego) following manufacturer instructions. CRISPR/Cas9 deletions were confirmed using Sanger sequencing of the amplified genomic region of interest and Inference of CRISPR Edits analyses (Synthego). Validation also involved immunoblotting to the cells with ferrochelatase and vinculin antibodies and PCR amplification to confirm the presence of the homozygous 107 bp out-of-frame deletion of the edited genomic region in K562-ALAS2 KO cells. *In vitro* erythropoiesis was monitored by following cellular differentiation of the cell lines (K562 and K562-ALAS2 KO) with benzidine staining and visualization on a disposable hemocytometer.

### 10X genomics single cell RNAseq and gene expression analysis

Blast cells from AML patient donors with over 60% blast expansion in marrow biopsies were isolated according to standard tissue-banking protocols, washed and resuspended at 10^6^ cells/ml. Cells were processed using the 10x Genomics Chromium controller and loaded according to standard protocol of the Chromium single cell 3’ library and v3 gel bead kit to capture 2,000 cells/Chromium Chip B. Single-cell capture, cell lysis, reverse transcription, and library preparation were performed per manufacturer instructions. Sequencing was performed on Illumina NextSeq 550 targeting150,000 reads/cell. The Cell Ranger Single-Cell software Suite (10x Genomics) was used for data processing, sample demultiplexing and gene expression quantification. t-Distributed stochastic neighbor embedding (t-SNE) and k-means clustering were used to reduce the data to a two-dimensional space and identify cell populations. Once the mean expression profiles of all cells were calculated, each cell was assigned to a subpopulation by the highest Spearman’s correlation.

### Gene expression analysis

The gene expression patterns related to heme biosynthesis from 10,000 patient-derived tumor samples were analyzed using the publicly available resources to study tissue-specific gene expression, the GTEx project (Ardlie; Lonsdale et al., 2013) and TCGA program (Chang et al., 2013).

### CRISPR pan-cancer gene essentiality analysis

The data from the DepMap Portal were used to calculate the gene essentiality scores in a similar manner to published methods published methods (Aguirre et al., 2016; Kim and Hart, 2021; Meyers et al., 2017). The gene essentiality score was used to evaluate the cell growth fitness. The lower the gene essentiality score value, the larger the gene loss effect on cell viability. The method as described in (Aguirre et al., 2016; Kim and Hart, 2021) as employed to correct the copy number bias in whole genome CRISPR/Cas screens by computing the mean of sgRNAs vs. the control plasmid library. For *in vivo* gene essentiality analysis in pancreatic and lung cancer models, the published data the published data(Zhu et al., 2021) were used to specifically examine the genes encoding the heme biosynthetic pathway enzymes and heme transporters.

### *In vitro* primary human hepatocytes

Prior to hepatocyte seeding, sterilized 384-well plates were coated with collagen and kept in a cell incubator at 37 °C overnight. After washing the wells, hepatocytes were suspended in supplemented growth medium, and the cell viability was subsequently quantified using the trypan blue exclusion method. The hepatocyte density was set at 1 × 10^3^ live cells μL^-1^, 18 μL cell suspension was added to each well and the medium was exchanged thrice weekly. Cells were incubated with 1.0 mM ALA at 37 °C for 4 hours. Imaging of live cells (ALA-treated and control) was performed upon Hoechst 33342 staining and using a CellInsight CX7 High-Content Screening Platform.

### *In vitro* hepatocyte cell line HC-04

After suspension of HC-04 hepatocyte cells and transfer to a collagen-coated T75 flask, the cells grew in a defined medium at 37 °C until reaching 70% confluence. Cells were trypsinized, washed with culture medium and seeded in 384-well plates at a density of 6000 cells/well in 20 μl of culture medium. Subsequent incubation with ALA, Hoechst 33342 staining and live cell imaging were as described for primary hepatocytes.

### Cellular PPIX quantification, TBHP-induced oxidative stress, and cell viability assay

Cellular PPIX accumulation was determined using fluorescence-activated cell sorting (FACS) as previously described (Fratz et al., 2014). PPIX accumulation in cells incubated in either the absence or presence of glycine or ALA was determined using fluorescence-activated cell sorting (FACS) as previously described. Upon tert-butyl hydroperoxide (TBHP)-induced oxidative stress and reactive oxygen species (ROS)-mediated erythroid differentiation, K562 and K562-ALAS2 KO cells were treated with 500 nM CellROX™ green reagent at 37 °C for 30 min. SYTOX™ Blue dead cell stain was uses to distinguish live (oxidatively stressed and nonstressed) from dead cells per manufacturer’s instructions. Cells were sorted by FACS, with the fluorescence emission of the SYTOX™ Blue-stained, dead cells being monitored at 440 nm (440/40 nm filter) upon excitation with an argon-ion 488 nm laser.

### Drug (“bait-and-kill”) assays

Viability and proliferation of HEL cells (ALA-treated and non-treated) were tested in the presence of either RSL3 (Yang et al., 2014) or ganetespib (Alexandrova et al., 2015) using using the CellTiter-Glo® luminescent reagent-based assay to determine cellular ATP as an indicator of metabolically active, and thus viable cells. The assays were performed according to manufacturer’s instructions, and the resulting luminescence of each assay was measured with a Clariostar Plus Microplate Reader (BMG Labtech).

### Additional methods and data are provided in supplemental data files

## Acknowledgments

We thank Charles Szerekes for facilitating FACS analysis and the USF genomics program for sharing computing resource and facilities. Also, we thank Rudwan M. Soukieh, Saeed Sinan and Thomas D. Nuhfer for discussions and for facilitating work related to this study.

RHYJ, GCF, GWR, and SMS acknowledge support from the Florida Department of Health Grant (BHC 9BC14). RHYJ also acknowledges support from ACS-Moffitt (ACS-IRG IRG-14-189-19), University of South Florida (Women’s Health Collaborative grant and the College of Public Health), and the Gates Foundation (OPP1023601 – for single-cell genomics protocol development)

## Author contributions

RHYJ, SRA, GCF, and GAH conceived the study. RHYJ, GCF, SRA, GAH, GWR, SMS and EMS devised the methodologies used for investigation. SRA, GAH, NEA, CM, and AB performed the mammalian cell and CRISPR experiments, and conducted *in vitro* biochemistry and biological function assays. RHYJ developed the computational algorithms and visualization tools used. RHYJ, GCF, GWR and SMS wrote the manuscript with contribution from SRA, and GAH. RHYJ, GCF, GWR, and SMS supervised the study.

## Data and materials availability

The data used in the study are deposited at XXX, and the raw data files are available at XXX. Supplementary Materials

## Materials and Methods

Figs. S1 to S8

Tables S1 to S8

