## Supplemental materials for "Porphyrin overdrive rewires pan-cancer cell metabolism"

**This Supplemental File Includes:**

**Materials and Methods**

**Supplementary Figures**

Fig. S1	Dependency on heme metabolism in many metastatic types of cancers

Fig. S2	*ALAS1* and *ALAS2* have different gene essentiality and pan-cancer gene expression patterns

Fig. S3	CRISPR/Cas9 gene targeting and inferred gene essentiality in cancer cells exemplify porphyrin overdrive

Fig. S4	Analysis of porphyrin overdrive in diverse cancer cell types

Fig. S5	*In vivo* CRISPR/Cas9 loss-of-function studies confirm essentiality of cancer porphyrin overdrive

Fig. S6	Analyses of gene expression in tumors of patients with diverse types of cancer support cancer porphyrin overdrive

Fig. S7	Cancer progenitor cells highly express cell niche interaction genes and show features of highly dynamic metabolic substrate trafficking hubs

Fig. S8	AML patient cancer progenitor cells show enhanced metabolic flux

**Other Supplementary Materials for this Manuscript Include:**

Supplementary Table S1	CRISPR/Cas9 essentiality screens related to heme metabolism and oncogenesis

Supplementary Table S2	CRISPR/Cas9 essentiality scores of diverse cell lines related to heme biosynthesis genes

Supplementary Table S3	Partial heme biosynthetic pathway requirement for *in vitro* cancer cell growth

Supplementary Table S4	*In vivo* murine CRISPR/CAS9 screen gene essentiality scores in pancreatic and lung cancers

Supplementary Table S5	Tissue-specific gene expression in patient-derived tumor samples (GTE_X_ project and TGCA program) *vs*. normal individual samples

Supplementary Table S6	Singe cell RNAseq of human embryonic stem cells

Supplementary Table S7	Single cell RNAseq of bone marrow samples from AML patients. Marker genes for the clusters of cell populations

Supplementary Table S8	Analysis of single cell RNAseq data for the bone marrow samples from AML patients using t-SNE
**Materials and Methods**

**Cell culture**

The human chronic myelogenous leukemia cell line K562 (ATCC, Cat No. CCL-243) and human erythroleukemia cell line HEL 92.1.7 (ATCC, Cat No. TIB-180) were obtained from the American Type Culture Collection. Both the cells were grown in RPMI 1640 medium (Gibco) supplemented with 10% fetal bovine serum (Sigma), gentamicin (1000x, Fisher, Cat No. 15-710-072), PennStrepNeo solution (100x, Fisher, Cat No. 15640-055) and 2 mM L-glutamine at 37 °C in a humidified 5% CO_2_ atmosphere. 5-Aminolevulinic acid hydrochloride (ALA), purchased from Alfa Aesar (Cat No. A16942ME), was dissolved in distilled water to yield a stock concentration of 1.0 M, and stored at -20 °C. Glycine, purchased from Fisher Chemical (Cat No. BP381-500), was dissolved in phenol red-free culture medium purchased from Gibco (Cat No. 11835055) to give a stock concentration of 1 M.

**Cell growth and quantification**

K562 or HEL cells were plated at 1-2x10^5^/mL, and cells were counted by trypan blue (Corning, Cat No. 25-900-CI) exclusion over time, with passage back to the starting cell density as needed. Total cell numbers were determined based on the passage dilution at each time point, on a disposable hemocytometer (Incyto, Cat No. DHC-N01).

**Generation of knockout cell lines by CRISPR**

K562 cells containing deletions in the ALAS2 or FECH genes were created using a multi-guide strategy via nucleofection (Lonza) of Cas9/RNP complexes (Gene Knockout Kit v2, Synthego) following manufacturer instructions. The guide RNAs utilized were: ALAS2: UGAAGGCUUUCAAGACAGGU, CAAUCUUGCUCUUCCCAUCC, and AGAUUCUCCAUCUUGGGCGA; FECH: UUAGACUCAUACCUCUUCUG, CUGGGCUGUUUCUGUGGUGA, and CUGACAGACCCUCCAGCUGC. CRISPR/Cas9 deletions were confirmed following amplification of the genomic region of interest, Sanger sequencing of the amplified region, and Inference of CRISPR Edits (ICE) analyses (Synthego). Cells were lysed in RIPA buffer with protease and phosphatase inhibitors and immunoblotted with antibodies that recognize ferrochelatase (SC-377377) and vinculin (SC-73614) (Santa Cruz Biotechnology). PCR amplification of first strand cDNA from K562 and K562-ALAS2 KO was used to confirm the presence of the homozygous 107 bp out-of-frame deletion that was detected in the ICE analyses of the edited genomic region in K562-ALAS2 KO cells.

***In vitro* erythropoiesis protocols**

*In vitro* erythropoiesis was monitored by following cellular differentiation of the cell lines (K562 and K562-ALAS2 KO) with benzidine staining. The benzidine solution was prepared by mixing 5 mL of 30% hydrogen peroxide (H_2_O_2_) with 1 mL of 0.2% benzidine dihydrochloride in 0.5 M acetic acid. Around one million cells were washed thrice with in Dulbecco's phosphate-buffered saline (DPBS, 1X; Corning, Cat No. 21-031-CV) before being resuspended in 250 μL of 1x DPBS mixed with 250 μL benzidine solution and incubated at room temperature for 10 minutes. The cells stained brown-blue were visually recognized and counted as positives, on a disposable hemocytometer (Incyto, Cat No. DHC-N01). The experiments were conducted in triplicates, and multiple microscopic fields were counted.

**AML patient single cell RNAseq and heme biosynthetic pathway-related gene expression analysis**

Cells were obtained from human donors with over 60% blast expansions in marrow biopsies. Blast cells were washed and isolated according to standard tissue-banking protocols. Cells were carefully washed in in Dulbecco's phosphate-buffered saline (DPBS, 1X; Corning, Cat No. 21-031-CV) and resuspended at 10^6^ cells/mL to avoid cell aggregates. Cells were processed using the 10x Genomics Chromium controller and the Chromium single cell 3’ library and gel bead kit (10x Genomics, Cat No. PN-1000075) following the standard manufacturer's protocols. First, gel beads-in-emulsion (GEMs) were generated by combining barcoded single cell 3’ v3 gel beads, a master mix containing cells, and partitioning oil onto Chromium chip B. To achieve single cell resolution, cells were delivered at a limiting dilution, such that the majority (~90-99%) of generated GEMs contain no cells, while the remainder contain predominantly a single cell. Between 2,000 - 21,000 live cells were loaded onto the Chromium controller to recover between 1,500 - 15,000 cells for library preparation and sequencing. Immediately following GEM generation, the gel beads were dissolved, primers were released, and any co-partitioned cells were lysed. An Illumina TruSeq Read 1 (read 1 sequencing primer), 16 nt 10x barcode, 12 nt unique molecular identifier (UMI) and 30 nt poly(dT) sequence were mixed with the cell lysate and a master mix containing reverse transcription (RT) reagents. Incubation of the GEMs produces barcoded, full-length cDNAs from poly-adenylated mRNAs. Next, GEMs were broken, and cDNA was amplified and quantified using an Agilent high sensitivity DNA screentape (Agilent Technologies, Cat No. 5067-5592). SPRIselect magnetic beads (Beckman Coulter, Cat No. B23317) were used to purify the first-strand cDNA from the post GEM-RT reaction mixture, which included leftover biochemical reagents and primers. Barcoded, full-length cDNA was amplified via PCR to generate sufficient mass for library construction. Enzymatic fragmentation and size selection were used to optimize the cDNA amplicon size. TruSeq Read 1 (read 1 primer sequence) was added to the cDNAs during GEM incubation. P5, P7, a sample index, and TruSeq Read 2 (read 2 primer sequence) were added via end repair, A-tailing, adaptor ligation, and PCR. The final library quality was assessed using an Agilent Bioanalyzer high sensitivity chip. Samples were then sequenced on the Illumina NextSeq 550 with a target of 150,000 reads/cell.

The Cell Ranger Single-Cell software Suite (10x Genomics) was used for data processing, sample demultiplexing and gene expression quantification. For data analysis, genes with more than one unique molecular identifier (UMI) counts were used. The top 1000 most variably expressed genes were used for further clustering, and t-Distributed Stochastic Neighbor Embedding (t-SNE) analysis was performed to reduce the data dimension to a two-dimensional space, and k-means clustering was used to identify cell populations. The mean expression of genes in all cells in a given cluster was calculated, and the expression of each gene was compared with that of the same gene in all the other clusters. For cell classification, the mean expression profiles of all cells were first calculated, and then each cell was assigned to a subpopulation by the highest Spearman’s correlation.

**Heme biosynthetic pathway-related gene expression analysis**

The gene expression patterns related to heme biosynthesis from patient-derived tumor samples were analyzed using the publicly available resources to study tissue-specific gene expression, the GTEx project (Ardlie; Lonsdale et al., 2013) and TCGA program (Chang et al., 2013). The heme biosynthetic pathway gene expression patterns in 10,000 tumors *vs*. matched normal tissues were used for analysis.

**CRISPR pan-cancer gene essentiality analysis**

The data from the DepMap Portal were used to calculate the essentiality scores in a similar manner to published methods (Aguirre et al., 2016; Kim and Hart, 2021; Meyers et al., 2017). The whole genome CRISPR/Cas9 datasets were used to identify significantly depleted mutant cells bearing a specific gene knock out in a pooled experiment. Gene essentiality was estimated from a given gene dependency inferred from CRISPR/Cas9 gRNA gene knockout. The essential score was used to evaluate the cell growth fitness. The lower the essentiality score value, the larger the gene loss effect on cell viability**.** Thus, a score of 0, < 0 and > 0 indicates no fitness change, fitness loss and fitness gain (*i.e*., possible growth advantage for the cell line) under the assay conditions, respectively). The method as described in (Aguirre et al., 2016; Kim and Hart, 2021) was employed to correct the copy number bias in whole genome CRISPR/Cas screens by computing the mean of sgRNAs *vs*. the control plasmid library. Commonly essential genes were required for the fitness of most cell lines across cancer types (Dempster et al., 2019; Pacini et al., 2021). For in *vivo* gene essentiality analysis in pancreatic and lung cancer models, the published data in (Zhu et al., 2021) were used to specifically examine the genes encoding the heme biosynthetic pathway enzymes and heme transporters.

***In vitro* primary human hepatocyte culture, quantification, and imaging**

Sterilized 384-well plates (Greiner, Cat No. 781091) were unpackaged in a class II biosafety cabinet and placed in a secondary container (*i.e.*, plates were placed in large assay pans) to serve as a lid and control evaporation. The day prior to hepatocyte seeding, wells were collagen coated with 40 μL of 15 μg μL^-1^ rat tail collagen I (Corning, Cat No. 354236) in sterile filtered 0.02 M acetic acid (Thermo Fisher Scientific, Cat No.), and kept at 37 °C overnight. Immediately prior to seeding the wells were washed thrice with sterile phosphate buffered saline (PBS) and then filled with 20 μL *in vitro* GRO® CP plate medium (BioIVT, Cat No. Z99029) supplemented with 1x Pen-Strep-Neo solution (100x, Fisher, Cat No. 15640-055) and 20 μM gentamicin (1000x, Fisher, Cat No. 15-710-072). Vials of cryopreserved (male) primary human hepatocytes (BioIVT, Cat No. M00995-P) were thawed by immersion in a 37 °C water bath for 2 minutes, sterilized with 70% ethanol in a sterile field, and the contents were added directly to 4 mL plate medium. Live and dead cells were quantified by trypan blue exclusion on a Neubauer improved hemocytometer. The hepatocyte density was set at 1 × 10^3^ live cells μL^-1^, and 18 μL cell suspension was added to each well. Medium was exchanged with the GRO® CP plating medium, described above, thrice weekly. The cells were incubated with 1.0 mM ALA at 37 °C for 4 hours. Both ALA treated and non-treated cells were handled under very low light conditions. During the last 45 min of incubation, a staining solution diluted in phenol-free, serum-free RPMI (Gibco, Cat No. 11835055) containing Hoechst 33342 (Life Technologies, Cat No. H3570) at a final concentration of 10 μM, was added to the cells. Live cell imaging was performed on a CellInsight CX7 High-Content Screening Platform (Thermo Fisher Scientific), and each plate well was counted for hepatocytes nuclei staining.

***In vitro* hepatocyte cell line HC-04 culture, quantification, and imaging**

Cryopreserved HC-04 hepatocyte cells were thawed, suspended into a previously prepared hepatocyte culture medium, and transferred to a T75 flask coated with collagen (Corning, Cat No. 354236) at 5 μg/cm^2^. The previously prepared hepatocyte cell-line culture medium consisted of a mixture of F12 base medium (Invitrogen, Cat No. 11765-054) and MEM base medium (Invitrogen, Cat No. A10490-01) on 1:1 (v/v) ratio, containing 10% FBS (Hyclone, Cat No. SH30070), 1.0 M HEPES (Invitrogen, Cat No. 15630-080) and200 mM glutamine (Invitrogen, Cat No. 25030-081) (Sattabongkot et al., 2006). Cells were allowed to grow until reaching 70% confluence, and the medium was changed every other day. Then, the cells were trypsin-hydrolyzed with TrypLE™ Express Enzyme (1X) (Gibco, Cat No. 12605028) and washed with hepatocyte culture medium. The HC-04 cells were seeded at a density of 6000 cells/well and cultured in 384-well plates (Greiner, Cat No. 781091) in 20 μl of the above medium/well. Cells were incubated either in the absence or presence of 1.0 mM ALA at 37 ^o^C for 4 hours. Both ALA treated and non-treated cells were handled under very low light conditions. During the last 45 min of incubation, a staining solution diluted in phenol-free, serum-free RPMI (Gibco, Cat No. 11835055) containing Hoechst 33342 (Life Technologies, Cat No. H3570) at a final concentration of 10 μM, was added to the cells. Live cell imaging was performed on a CellInsight CX7 High-Content Screening Platform (Thermo Fisher Scientific), and each plate well covering 15 fields at 20x was counted for hepatocytes nuclei staining.

**Cellular PPIX quantification**

Protoporphyrin (PPIX) accumulation was determined using fluorescence-activated cell sorting (FACS) as previously described (Fratz et al., 2014). K562 and K562-ALAS2 KO cells were independently seeded in 24-well plates for suspension culture (Greiner, Cat No. 662102) overnight at a density of 1.2 x 10^5^ cells/well. The K562 and K562-ALAS2 KO cells were incubated in medium (described in the Cell Culture section) containing 1.0 mM ALA, at 37 °C for 4 hours. Preparation of either 5-ALA-treated cells or control (non-ALA-treated) cells was under very low light conditions. Following incubation, cells were washed thrice with Dulbecco's phosphate-buffered saline (DPBS, 1X, Ca^2+^- and Mg^2+^-free; Corning, Cat No. 21-031-CV) and resuspended in 250 μL of 1x DPBS. Briefly, cells were washed once with serum-free medium (Gibco, Cat No. 11835055) and incubated in 6-well plates containing the medium (described in the Cell Culture section) in either the absence or presence of 100 mM glycine or ALA (0.1 mM, 0.25 mM, 0.5 mM, and 1.0 mM) at 37 °C. Intracellular PPIX concentration was measured 18 hours later by FACS. The cell suspension was passed through a 40-μm Flowmi™ Cell Strainer to eliminate clumps and debris prior to transferring to BD Falcon tubes under very low light conditions to minimize phototoxicity caused by PPIX accumulation. FACS analyses were performed using a BD LSR II Analyzer (Becton, Dickinson, and Company) and FACSDiva Version 6.1.3 software. To eliminate any background red fluorescence, the 633 nm-red laser was blocked during the collection of the PPIX emission data. PPIX emission was determined in the 619 nm and 641 nm range (630/22BP filter) upon excitation of the cells with the 405 nm laser. Forward-scatter (FSC) versus side-scatter (SSC) dot plots were used to gate the whole cells and thus remove the contribution of the cell debris from the population being examined. A minimum of 10,000 of the gated cells was then depicted in dot plots of SSC *vs*. PPIX fluorescence; the gate was defined based on wild-type K562 cells without any perturbation as negative controls.

**Cellular reactive oxygen species (ROS) detection and cell viability assays**

Both K562 and K562-ALAS2 KO cultures were seeded in 24-well plates for suspension culture (Greiner, Cat No. 662102) at a density of 1.2 x 10^5^ cells/well in the medium described in the Cell Culture section, at 37 °C overnight. After the incubation, cells were washed thrice with Dulbecco's phosphate-buffered saline (DPBS, 1X, Ca^2+^- and Mg^2+^-free; Corning, Cat No. 21-031-CV) and resuspended in 250 μL of 1x DPBS and stained for 30 min at 37 °C with 500 nM CellROX green reagent (Life Technologies, Cat No. C10444). During the last 15 min of staining, 1 μl of the 5 μM SYTOX Blue Dead Cell stain (Life Technologies, Cat No. S34857) was added to distinguish live from dead cells. Cells were immediately analyzed by fluorescence-activated cell sorting (FACS) using a BD LSR II Analyzer (Becton, Dickinson, and Company) and FACSDiva Version 6.1.3 software. Emission of the oxidized fluorogenic CellROX™ green was measured at 525 nm (530/30 BP filter) upon excitation of the cells using the 488 nm laser. K562 cells treated with 250 μM tert-butyl hydroperoxide (TBHP, Thermo Scientific, Cat No. 180340050) for 15 min were used as a positive control. K562 cells treated with 1 mM ALA for 4 hours as “PPIX fluorescence positive control”. Both ALA-treated and non-treated cells are handled under very low light conditions.

**Drug (“bait-and-kill”) assays**

The compounds RSL3 (MedChemExpress, Cat No. HY-100218A) and ganetespib (MedChemExpress, Cat No. HY-15205) were dissolved in 100% DSMO (ATCC, Cat No. 4-X) to yield 10 mM stocks, which were stored at −80 °C until further use. HEL cells were seeded in 38-well plates (Greiner, Cat No. 781091) at a density of 6000 cells/well and in 20 μl of medium (described in the Cell Culture section) per well. The cells were allowed to proliferate for the next 48 hours. Both ALA treated and non-treated cells were handled under very low light conditions throughout the assay. The cells were treated with 1 mM ALA in individual plates for 4 hours and 24 hours. Both ALA-treated and non-treated plates were tested with the compounds RSL3 and Ganetespib in triplicate wells using an 18-point concentration format with 2-fold dilutions (final concentrations of 132 µM to 1 nM) bringing the total volume to 25.6 μl per well. Cell proliferation was measured utilizing the CellTiter-Glo 2.0 reagent (Promega, Cat No. G9243) to quantify cellular ATP according to manufacturer's instructions by adding 25.6 μl of the reagent per well. Luminescence was measured with a Clariostar Plus Microplate Reader (BMG Labtech). For each assay plate, a DMSO control (0.1%), a positive control, a negative control and blanks were added, and a minimum of 12 wells per plate were analyzed. Data were reported as arbitrary luminometric units (ALU).

(Socolovsky, 2013)

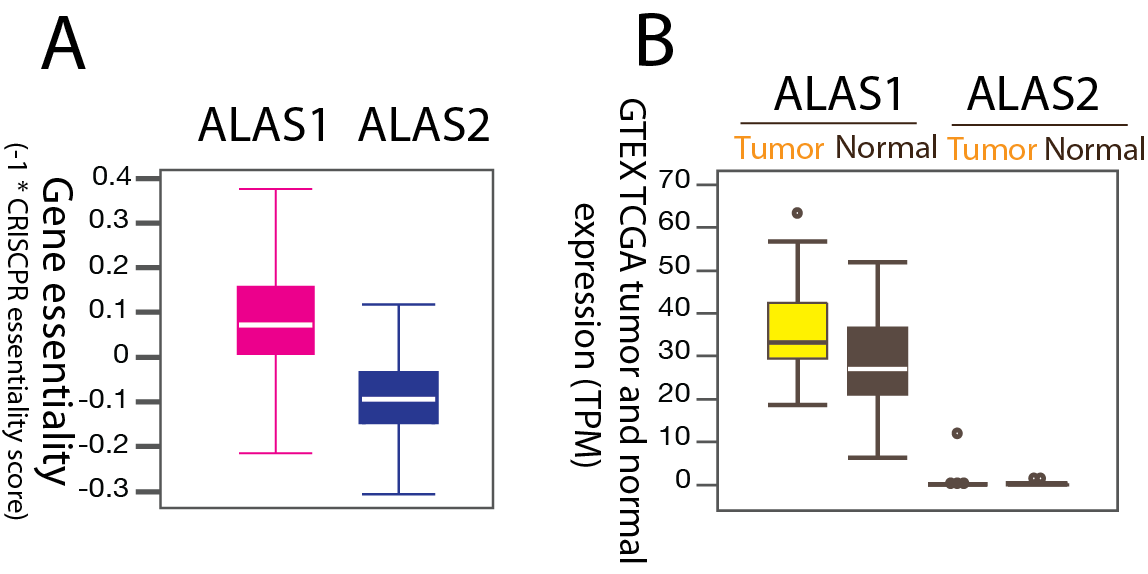

**Supplementary Figure S1.** **Dependency on heme metabolism in many metastatic types of cancer.** The data for the CRISPR/Cas9-gene targeting of the genome-scale loss-of-function screens in a set of cancer cell lines were retrieved from DepMap. The columns refer to different cancer cell lines, while the rows refer to specific genes. Note the differences between the loss-of-function scores for specific proteins associated with heme biosynthesis (UROD and FECH) in this panel of metastatic cancer cell lines. [Abbreviations: AURKA, Aurora kinase A; AURKB, Aurora kinase B; CYC1, cytochrome c-1; DGCR8, DiGeorge syndrome critical region 8; FECH, ferrochelatase; HMOX1, heme oxygenase 1; HPX, hemopexin; SDHC, succinate dehydrogenase complex subunit C; TFRC, transferrin receptor; UROD, as uroporphyrinogen III decarboxylase]

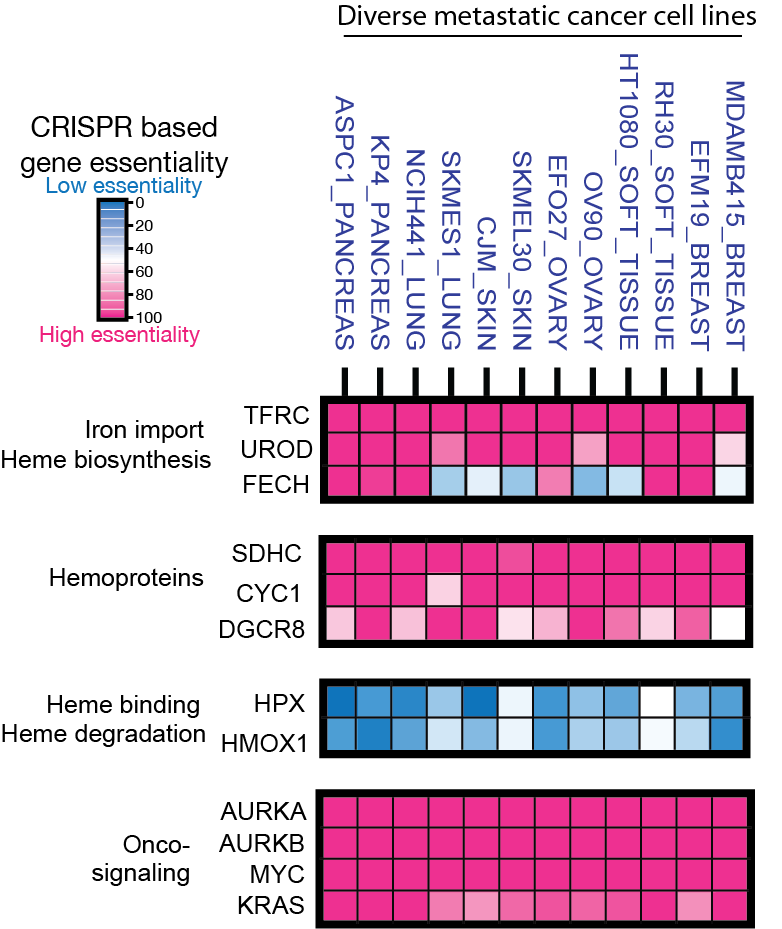

**Supplementary Figure S2. ALAS1 and ALAS2 have different gene essentiality and pan-cancer gene expression patterns.** (**A**) The isozyme gene pair *ALAS1* and *ALAS2* have different gene essentiality based on CRISPR/Cas9 loss-of-function results. ALAS2 was found not to be essential in any of the 27 major cancer types in over 300 cell lines, including erythroleukemia cell lines. The *Y*-axis label “CRISPR KO gene dependency” refers to the essentiality of *ALAS1* or *ALAS2*. The essential score was used to evaluate the cell growth fitness. An essentiality score of 0, < 0 and > 0 indicates no fitness change, fitness loss and fitness gain under the same experimental conditions, respectively. (**B**) A study of ~10,000 patient derived tumors from GTEx and TCGA shows that *ALAS1* expression is elevated in tumors compared to normal cells, while like in normal cells, *ALAS2* expression is absent in most cells, except myeloid leukemia cells. However, the *ALAS2* expression levels in normal erythroblasts do not significantly differ from those in cancer erythroblasts and myeloid leukemia cells. The dots indicate outliers of the samples with respect to *ALAS* expression. TPM, transcript per million.

**Supplementary Figure S3.** CRISPR/Cas9 gene targeting and inferred gene essentiality in cancer cells exemplifies porphyrin overdrive. A total of over 300 cancer cell lines from 27 major cancer types are analyzed. The data set represents 18 human tissue types that give rise to diverse groups of cancer. Whole genome loss-of-function growth phenotypes show most cancer cells only require a partial heme biosynthetic pathway to survive. Gene essentiality is estimated from gene X dependency inferred from CRISPR/Cas9 gRNA gene X knockout. Low and high cell viability with deletion of gene X indicate that the cancer cells have a high and low dependence on gene X for survival, respectively. Thus, the lowest and highest gene essentiality values are associated with the least and most profound dependence of the cells on the loss of gene X-dependent, respectively. The UROD gene for the fifth enzyme of the pathway has the highest essentiality in all types of cancers, while genes encoding other enzymes are dispensable in many cancer cell lines. Each of the rows in the 18 human tissue panels represents a distinct cancer cell line for the pertinent tissue. [ABCG2, ATP-binding cassette (ABC) transporter subfamily G, member 2; ALAD, ALA dehydratase (aka porphobilinogen synthase); ALAS1, 5-aminolevulinate synthase 1; CPOX, coproporphyrinogen oxidase; FECH, ferrochelatase; FLVCR, feline leukemia virus subgroup C receptor family; HMBS, hydroxymethylbilane synthase; PPOX, protoporphyrinogen oxidase; SLC48A1, solute carrier family 48, member 1, a.k.a heme transporter HRG1; UROD, uroporphyrinogen III decarboxylase; UROS, uroporphyrinogen III synthase]

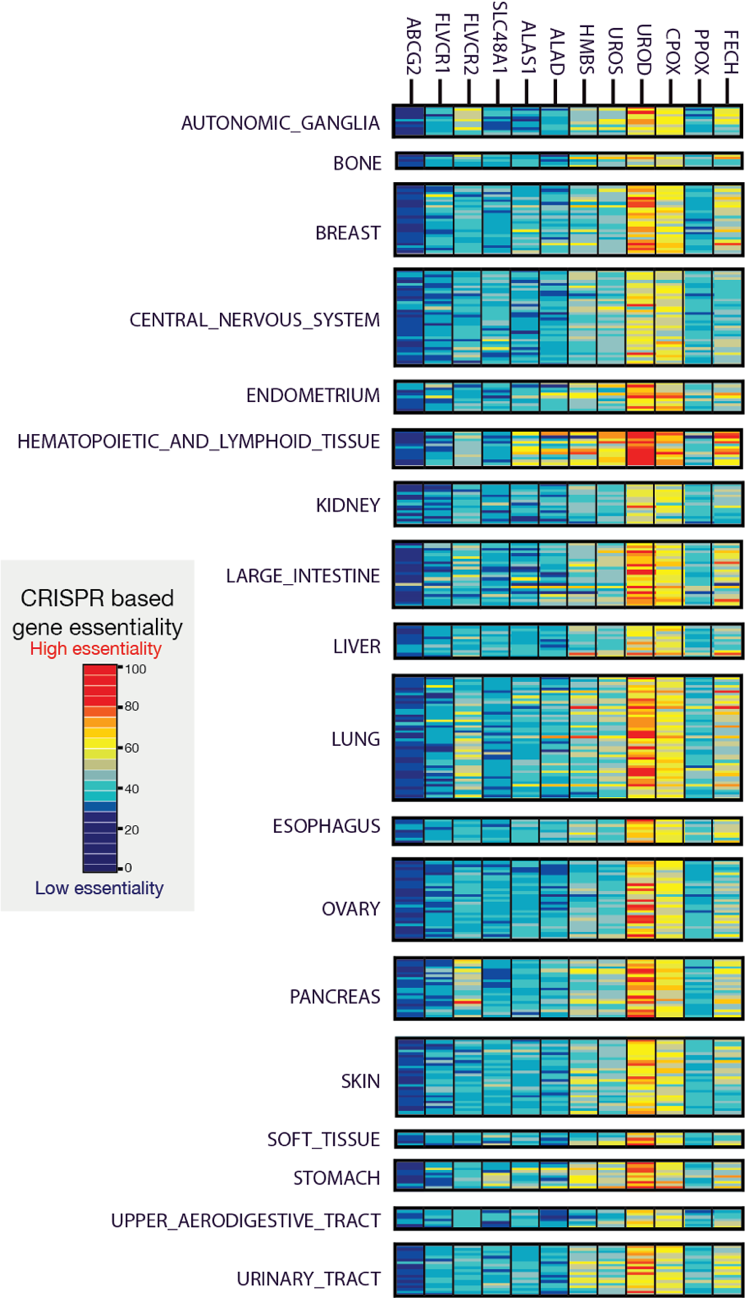

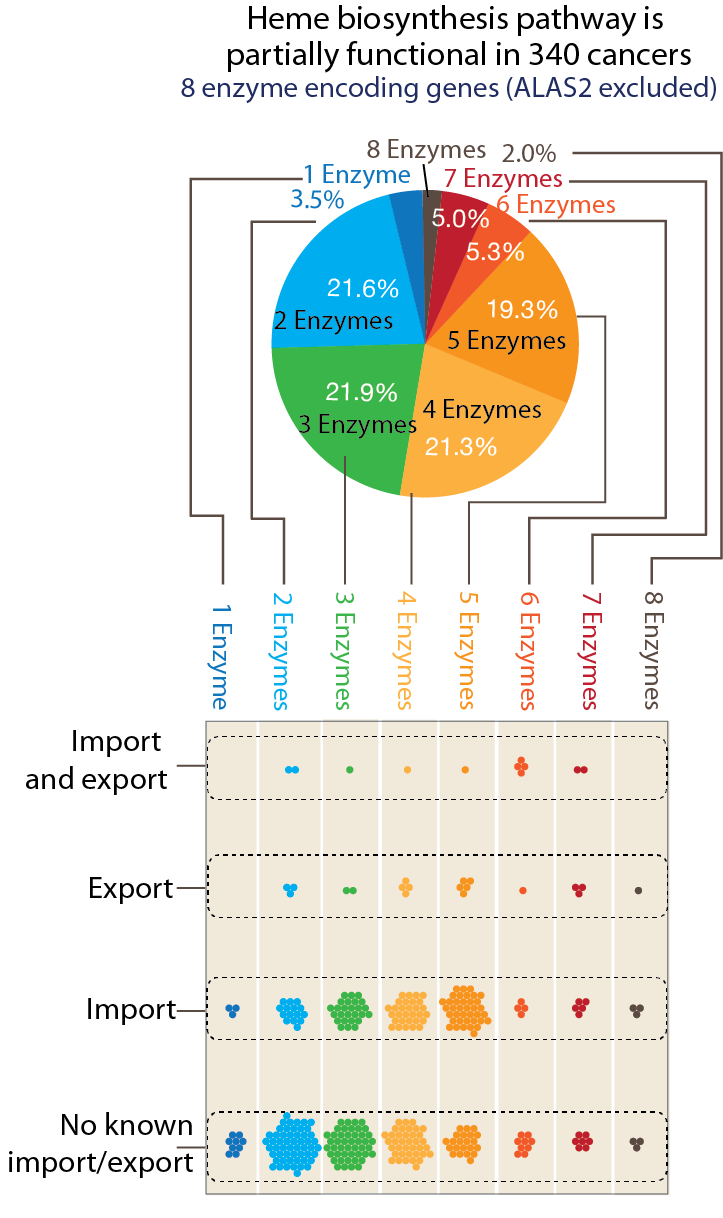

**Supplementary Figure S4. Analysis of porphyrin overdrive in diverse cancer cell types**. Most cancer cells from diverse lineages (27 major cancer types) representing 18 tissues show features of porphyrin overdrive based on CRISPR KO gene essentiality analysis. There are 8 enzymatic steps for heme biosynthesis, represented by nine genes with 2 different genes for the first enzyme ALAS. *ALAS2* was not included in this analysis because it is an erythroid lineage-specific gene. Classification of porphyrin overdrive metabolism in eight groups is based on the number of essential enzyme-encoding genes in the heme biosynthetic pathway. Each of the groups is represented by a colored sector of the pie chart with an area corresponding to the percentage of the cancers with a specified partial number of functional heme biosynthesis enzymes as inferred from the gene essentiality results. For each of these groups, the gene essentiality of known heme importers (FLVCR2, SLC48A1) and exporters (FLVCR1, ABCG2) was also characterized. The number of trafficking genes is likely a conservative estimate as the field is still developing. Over 80% of the total cancer cells require only a “partial” heme biosynthetic pathway of 2-5 functional enzyme genes to survive, indicating the widespread ‘incomplete’ heme biosynthesis in cancer cells. One single enzymatic step suffices in 3.5% of all tested cancer cells, and the data reflect the heightened requirement of the intermediate enzyme/genes (*e.g.,* HBMS, UROS, UROD and CPOX).

**Supplementary Figure S5. *In vivo* CRISPR/Cas9 loss-of-function studies confirm essentiality of cancer porphyrin overdrive**. (**A**) Porphyrin overdrive, resulting from absence of heme metabolism hemostasis, is indicated by the essentiality of the genes for the intermediate enzymatic steps of heme biosynthesis in both murine pancreatic and lung cancer models. The figure shows the comparison of gene essentialities between *in vitro* and *in vivo* cancer models, and their statistically significant differences (column 3 of both panels) are indicated with *. The different shades of magenta indicate the different degrees of gene essentiality. The color scheme reflects the pancreatic essentiality in the study. Both *in vitro* and *in vivo* essentiality data were retrieved from Zhu *et al.* The increased gene essentiality, from *in vitro* to *in vivo*, of intermediate step of heme biosynthesis genes (HBMS, UROS, CPOX, PPOX) are the only shared metabolic essentiality between pancreatic and lung cancers are indicated with *. The *in vitro* gene essentiality data indicate that the functional intermediate enzymatic steps in heme biosynthesis are independent of tumor origin or tissue environment (Zhu et al 2021). (**B**) Analysis of ~2900 metabolic-related genes (retrieved from ref 9) show similar gene essentiality in the *in vitro* and *in vivo* settings, indicating that the survival of tumor cells depends on the examined encoded-metabolic enzymes both in vitro and in murine pancreatic and lung cancer models. [No statistically significant essentially differences were observed for the essentialities of *ALAS1*, *ALAS2*, and *FECH*, the genes for the first and terminal enzymes of the heme biosynthetic pathway, in either of the *in vivo* murine cancer models. [adj, adjusted.]

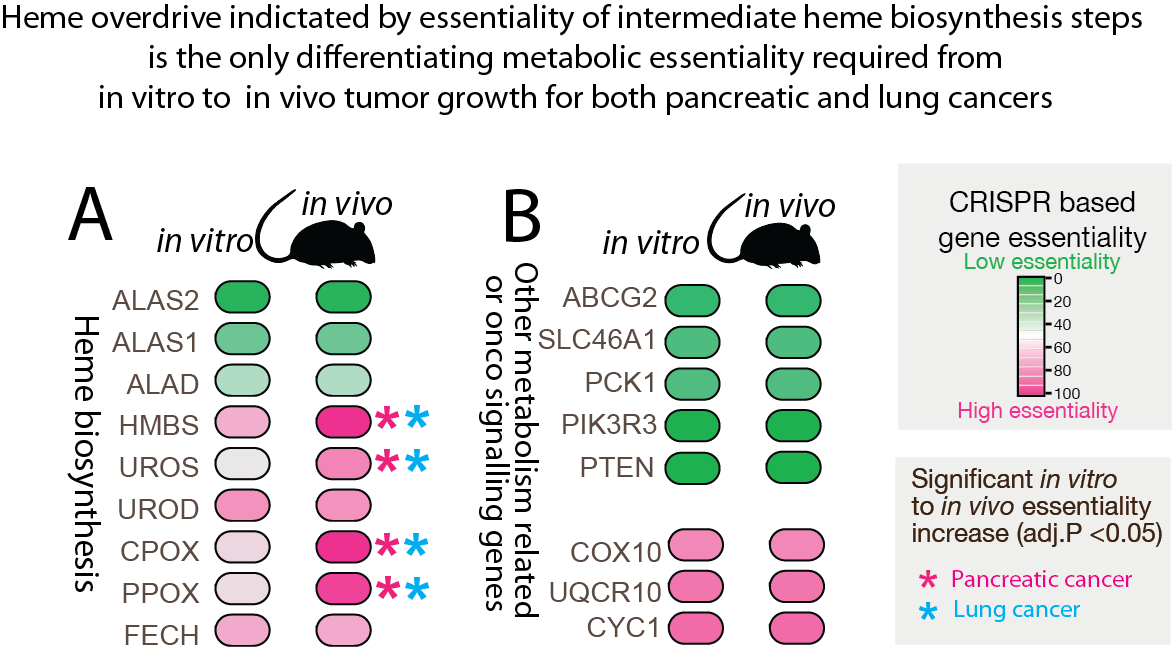

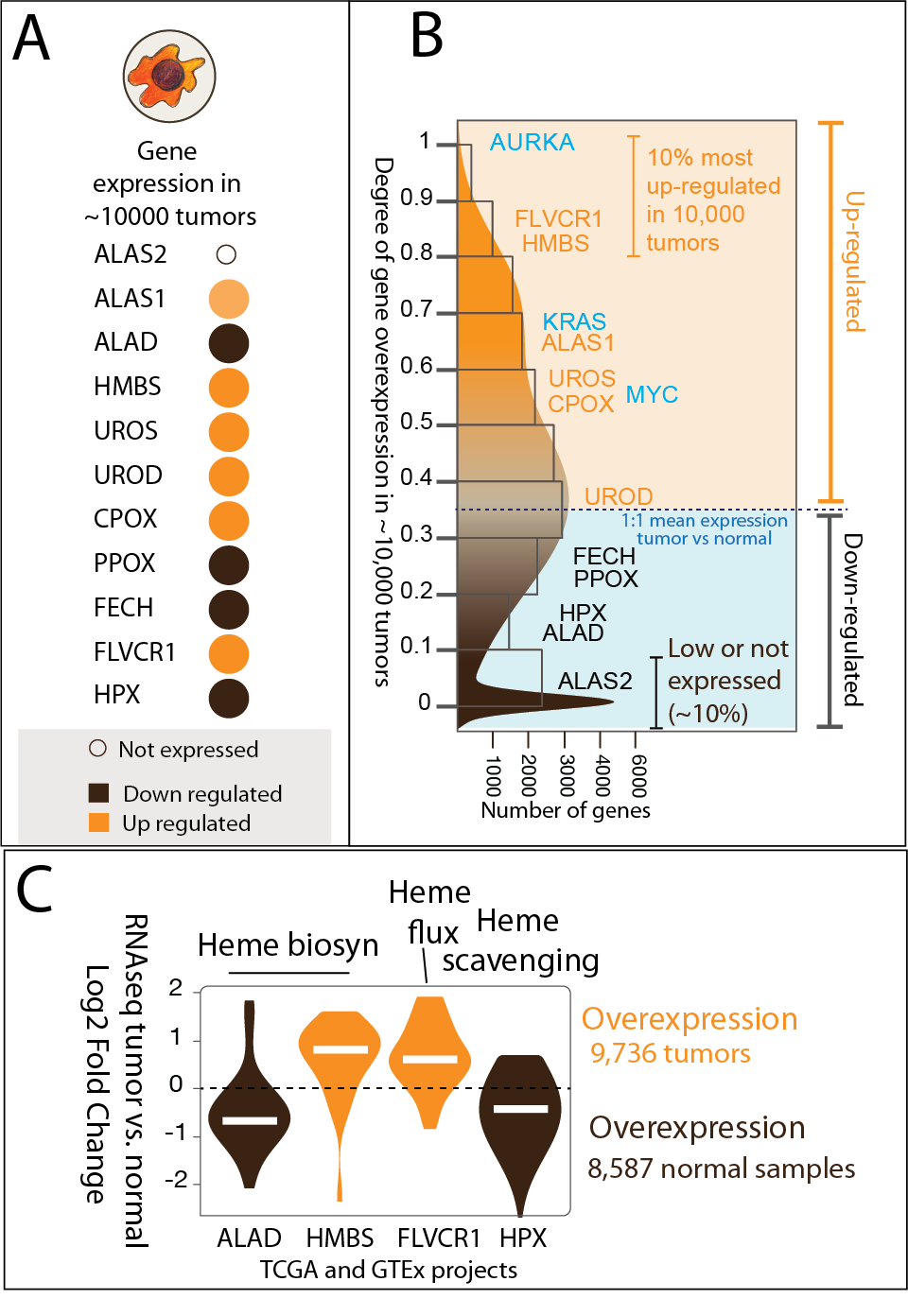

**Supplementary Figure S6. Analyses of gene expression in tumors of patients with diverse types of cancer support cancer porphyrin overdrive**. (**A**) Up-regulation of gene expression for some enzymes of the heme biosynthetic pathway in ~10,000 patient-derived tumors. *ALAS2* is only expressed in myeloid leukemias. (**B**) The genes for HMBS, the fourth enzyme of the heme biosynthetic pathway, and FLVCR1, the heme exporter, are among the most upregulated in over 80% of all tumors. Oncogenic signaling genes (in blue) *AURKA*, *KRAS* and *MYC* are upregulated in over 90%, 60% and 50% of all tumor types, respectively. Orange shading denotes upregulation of gene expression; blue shading denotes down-regulation of gene expression as compared to normal tissue. The scale of the *Y*-axis indicates the degree of over- or under- expression. (**C**) Expression of *ALAD*, encoding the second enzyme of the heme biosynthetic pathway, is downregulated in tumors, while the gene for the third enzyme of the pathway, HMBS, is overexpressed in most of the tumors. *FLVCR1*, encoding a heme exporter, is overexpressed in tumors, while *HPX*, for the heme scavenger hemopexin, is more abundantly expressed in normal tissues. Data are from the GTEx project and TCGA program. FC, fold change; scRNAseq, single-cell RNA sequencing.

**Supplementary Figure S6. Analyses of gene expression in tumors of patients with diverse types of cancer support cancer porphyrin overdrive**. (**A**) Up-regulation of gene expression for some enzymes of the heme biosynthetic pathway in ~10,000 patient-derived tumors. *ALAS2* is only expressed in myeloid leukemias. (**B**) The genes for HMBS, the fourth enzyme of the heme biosynthetic pathway, and FLVCR1, the heme exporter, are among the most upregulated in over 80% of all tumors. Oncogenic signaling genes (in blue) *AURKA*, *KRAS* and *MYC* are upregulated in over 90%, 60% and 50% of all tumor types, respectively. Orange shading denotes upregulation of gene expression; blue shading denotes down-regulation of gene expression as compared to normal tissue. The scale of the *Y*-axis indicates the degree of over- or under- expression. (**C**) Expression of *ALAD*, encoding the second enzyme of the heme biosynthetic pathway, is downregulated in tumors, while the gene for the third enzyme of the pathway, HMBS, is overexpressed in most of the tumors. *FLVCR1*, encoding a heme exporter, is overexpressed in tumors, while *HPX*, for the heme scavenger hemopexin, is more abundantly expressed in normal tissues. Data are from the GTEx project and TCGA program. FC, fold change; scRNAseq, single-cell RNA sequencing.

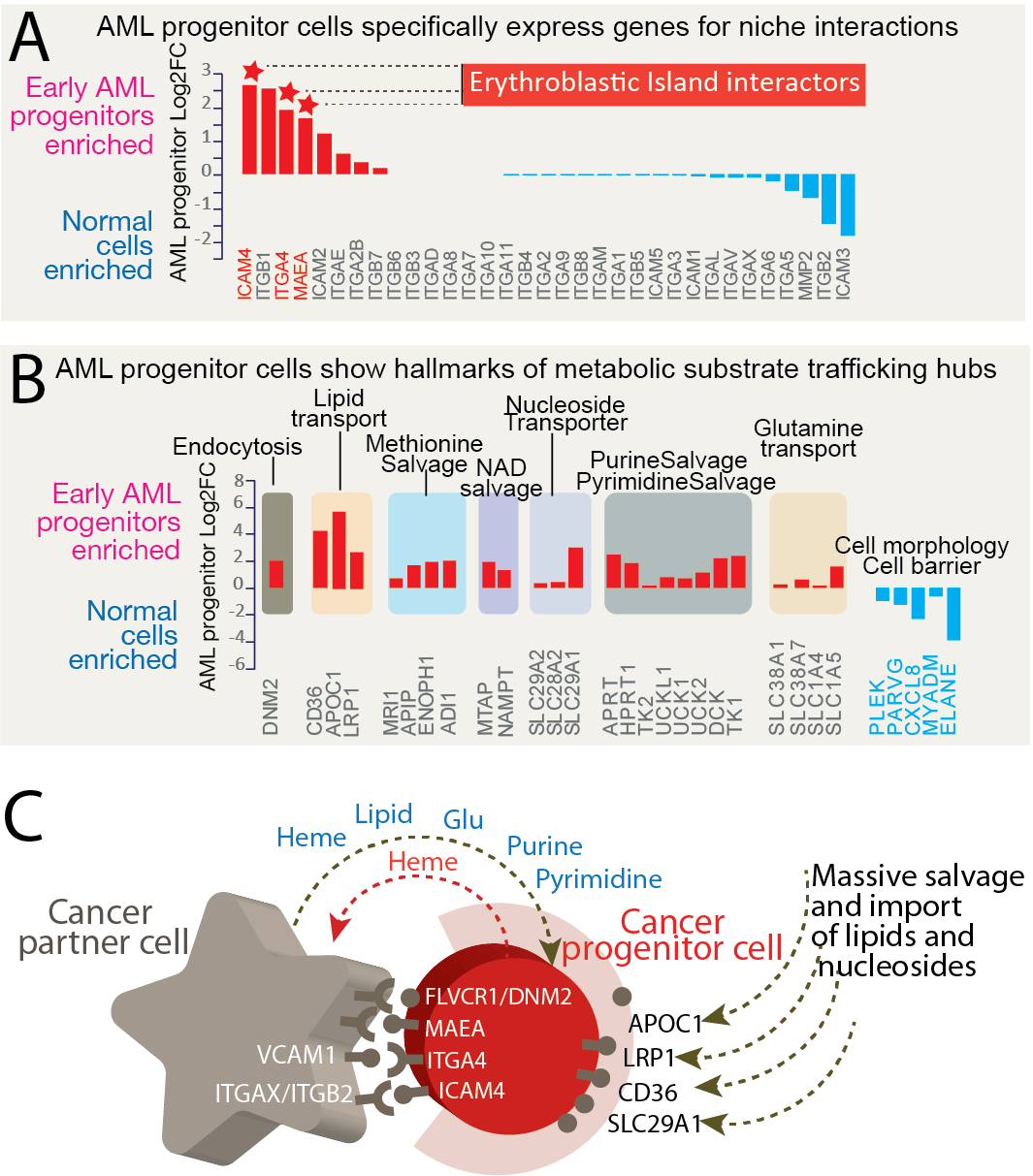

**Supplementary Figure S7. Cancer progenitor cells highly express cell niche interaction genes and show features of highly dynamic metabolic substrate trafficking hubs**. (**A**) AML progenitor cells specifically express genes for interacting proteins (“interactors”) within the erythroblastic island, the specialized bone marrow niche where erythroid precursors proliferate, differentiate, and enucleate. The *Y*-axis indicates the log_2_-fold change of gene expression levels in early AML progenitor cells relative to other progenitor cells. Positive log_2_-fold change values indicate increased expression levels over those of the control, while negative values indicated decreased expression relatives relative to those of the control. Names of selected genes are indicated in the *X*-axis. FC: Fold change. (**B**) AML progenitor cells show hallmarks of metabolic substrate trafficking and abundant salvaging pathways. Genes associated with endocytosis, lipid transport, NAD salvage, nucleotide salvage are upregulated, while genes related to cell morphology and barrier formation are downregulated. (**C**) The cancer cell niche is inferred to present elevated gene expression for proteins involved in intracellular surface molecular interactions (*e.g*., VCAM1-ITGA4 and ITGAX/ITGB2-ICAM4). The schematic represents a reconstruction of a cancer partner cell-cancer progenitor cell interaction based on previously established molecular interactions (Socolovsky 2013).

**Supplementary Figure S8. AML patient cancer progenitor cells show enhanced metabolic flux**. (**A**)The cancer early progenitor cells represent only 2% of the AML patient sample cell population (see Fig. 3). Light blue shading represents down-regulated genes and light pink shading indicates upregulated genes. The -log_10_ (p values) in the Volcano plot represent the level of significance of each gene, while the log_2_-fold change values represent the difference between the levels of expression for each gene between the AML progenitor cells and other populations of cells. The genes related to metabolic flux are highlighted in yellow and include genes associated with the postulated porphyrin overdrive, lipid import and macromolecule salvage. The canonical erythropoiesis master transcription regulators are colored as early (blue dots) and late (pink dots) stages based on their gene expression sequence during erythropoiesis. (**B**) The cancer progenitor specific gene expressions of APOC1 and S100A6 are shown in different cell populations. FPKM, fragments per kilobase million.

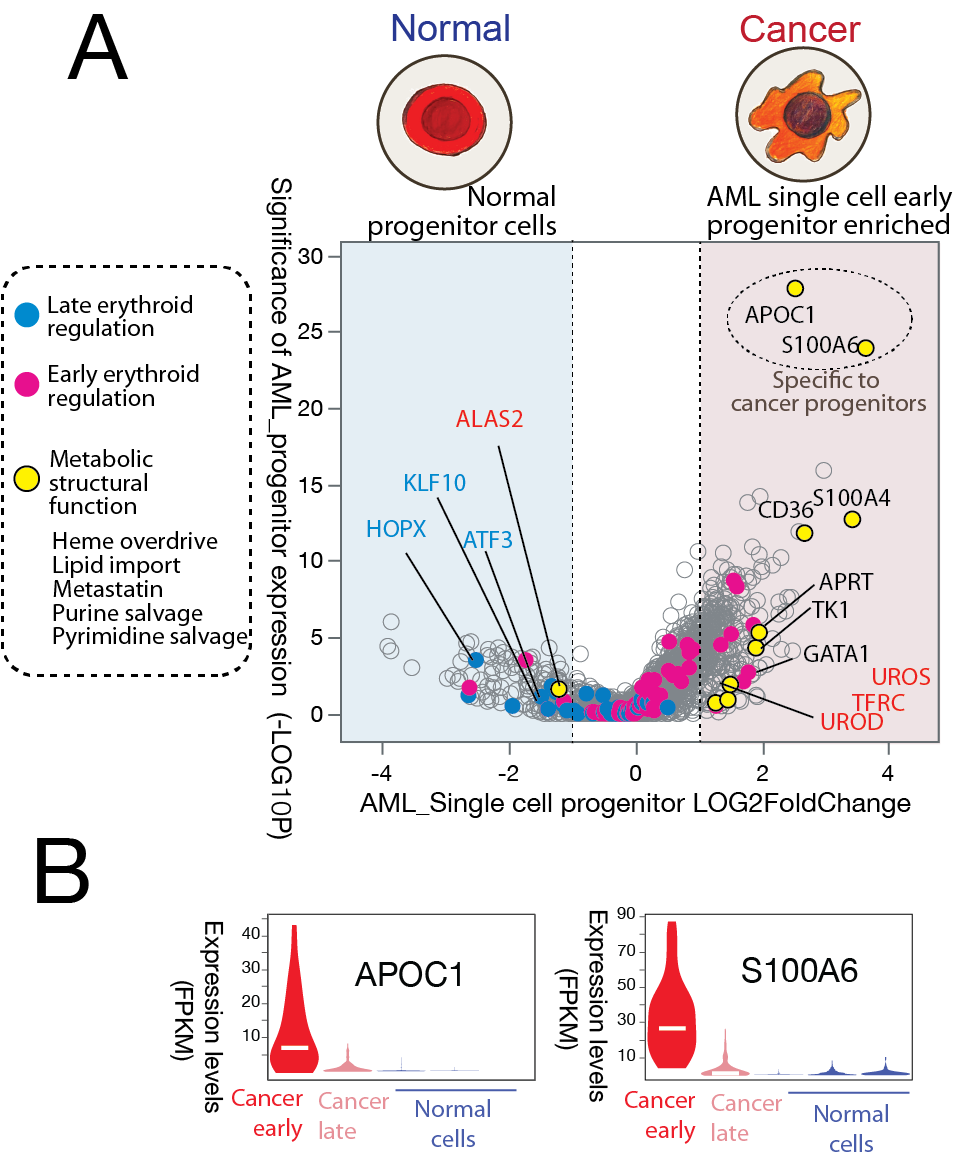
